# Probabilistic projections of distributions of kin over the life course

**DOI:** 10.1101/2025.03.24.644937

**Authors:** Joe Butterick, Jason Hilton, Jakub Bijak, Peter W. F. Smith, Erengul Dodd

## Abstract

**BACKGROUND:** Mathematical kinship demography is an expanding area of research. Recent papers have explored the expected number of kin a typical individual should experience. Despite the uncertainty of the future number and distributions of kin, just one paper investigates it.

**OBJECTIVE:** To develop a new method for obtaining the probability that a typical population member experiences one or more of some kin at any age through the life course.

**METHODS:** We use combinatorics and matrix algebra to construct and project a discrete probability distribution of kin. Our model requires as inputs, age-specific mortality and fertility.

**CONCLUSIONS:** We derive probabilities of kin-number for fixed age of kin and over all possible ages of kin. We derive expected numbers and variance of kin. We demonstrate how kinship structures are conditional on familial events.

**CONTRIBUTION:** The paper presents the first analytic approach allowing the projection of a full probability distribution of the number of kin of arbitrary type that a population member has over the life course.

## 1 Introduction

Family relationships are a fundamental part of the social structures of human societies (Alburez-Gutierrez et al., 2022). Kinship ties have impacts on physical and mental wellbeing (Vlachantoni et al., 2024), social and economic inequalities between and within generations (Verdery and Margolis, 2017; Margolis et al., 2024), and social and fiscal policy (Pittavino, Arpino, and Pirani, 2025). However, direct data on kinship relationships are rarely collected, and generally only available for limited samples or in countries with high quality demographic registers (Kolk, 2017). Studies that quantify the temporal changes in kinship networks are therefore extremely valuable.

In particular, understanding inequalities resulting from kinship relationships requires quantification of the diversity of kinship structures within the population. In addition, the future numbers and structures of kin remain uncertain. Micro-simulation facilitates the study of these kinds of structures, offering rich and detailed output about family composition both inside and outside the household (e.g. Wachter, Hammel, and Laslett, 1978; Hammel, 2005; Margolis and Verdery, 2019). However, micro-simulation is computationally expensive and technically challenging to set up. This can be a barrier to its deployment (notwithstanding creditable recent work in opening up access to these tools, such as Thiele et al., 2023). As such, analytical models of kinship relationships are highly desirable for their much greater computational efficiency.

Recent years have witnessed a resurgence in mathematical modelling of kinship (Caswell, 2019, 2020, 2022, 2024; Caswell and Song, 2021; Alburez-Gutierrez et al., 2021), providing, for example, richer insight into the future structures of kinship (Alburez-Gutierrez, Williams, and Caswell, 2023). Models of kinship have been applied to produce expected numbers of kin distributed by age or *stage* (or *state*) and sex for a representative population member, both within static and time-varying demographies. By stage, one refers to any population characteristic for which individuals can transition in and out of – a dimension out of scope for the present study. Despite the expanding interest in the estimation of kinship, only one model developed since 2019 has explored beyond the mean number of kin (see Caswell, 2024).

Population dynamics in the demographic context are driven by stochastic birth and death processes. That is, each (female) population member gives birth to a non-negative number of offspring (possibly zero), with variation between individuals’ reproductive profiles. Each population member either survives to the next age-class or dies, giving rise to variation in age-specific mortalities. Demographers usually refer to these probabilistic events as “demographic stochasticity” (Keyfitz and Caswell, 1997). Because population dynamics are governed by stochastic processes, and kinship dynamics are an emergent property of the population, an individual’s network of kin unfolds probabilistically.

This research complements and extends mathematical kinship demography by presenting a method to account for demographic stochasticity within the kinship network. Our paper is structured as follows: Section 1.1 provides a brief review of the relevant literature and existing mathematical frameworks for analysing kinship, while clarifying what sets this work apart. Section 1.2 outlines notation used. In Section 2 we introduce our assumptions and pre-requisites for formally constructing the model: Section 2.1 outlines the general demographic assumptions and Sections 2.2-2.6 lay the mathematical foundations. Section 3 provides our novel derivations, organised into analytically distinct regimes. In Section 4 we apply the framework using UK age-specific mortality and fertility data, sourced from the Human Mortality Database (2024) and Human Fertility Collection (2024). We illustrate the analysis by showing selected model output and explore its implications. We conclude in Section 5 by considering the contribution, prospects and limitations of this theoretical development with respect to the leading frameworks on kinship.

### 1.1 Relevant mathematical frameworks

There is a rich history of mathematical modelling of kinship which have considered demographic stochasticity. Already in the 1980s, research by Waugh (1981) and Joffe and Waugh (1982) applied branching processes with non-overlapping generations, namely the so-called Galton-Watson process, primarily focusing on biology and population ecology. Advancing on such methods Jagers (1982) and Jagers and Nerman (1984) included age structures (thereby accounting for overlapping generations) by utilising so-called “Crump-Mode-Jagers” (CMJ) branching process (Crump and Mode, 1968). Although very relevant to the field of kinship research, due to their complexity, these papers proved for the most part too technically complicated to allow their wider adoption.

In a recent remarkable paper, Caswell (2024) treats an individual’s kin as a population of their own, and subject to the same variability in births and deaths as any other population. The author’s proposed model accounts for demographic stochasticity through projecting the mean and variance of an unknown kin-number distribution. The matrix projection model proposed by Pollard (1966), emulating a multi-type Galton-Watson process, is applied to do so. Using the mean and variance in kin-number, Caswell (2024) subsequently assumes appropriate statistical distributions to represent kin, fits prediction intervals, and thereby estimates uncertainty associated with kin-number distributions.

While the proposed framework is more accessible to demographers, ecologists, mathematicians, and perhaps the occasional physicist, the technical modelling approach can be challenged on theoretical grounds. For instance, although the distribution of kin will always be discrete with a non-zero support on the non-negative integers, an appropriate choice might require specific knowledge. Caswell (2024) states that Poisson distributions should apply when the mean is close to the variance and cites previous work which adheres to this assumption (Caswell, Margolis, and Verdery, 2023). Moreover, previous research has shown that using a Galton-Watson process with age-structure coerced as “type” can under-estimate variation in population sizes (Mode, Jacobson, and Pickens, 1987).

An alternative approach would be to calculate the exact probability distribution of kin-number. To do so, one could apply similar methods to Tuljapurkar et al. (2020) who directly calculate the so-called lifetime reproductive success (i.e., a distribution for the number of offspring an individual has over their life course). The benefit of this method, emphasised by Tuljapurkar et al. (2020), is that by constructing a probability distribution for the number of offspring, one can account for variability in reproduction between individuals. Another obvious benefit is that higher-order moments become readily available.

In this research we develop a methodology for projecting the kin-number of a typical female population member in a one-sex time-invariant age-structured demography (see the illustration in Figure 1 for kin-types). Following the convention of the leading kinship models (Caswell, 2019, 2020; Caswell and Song, 2021; Caswell, 2022) we refer to this individual as ‘Focal’. We want to know Focal’s kin-network at each age of Focal’s life. In the same vein as Caswell (2024), we seek to account for demographic stochasticity in the kinship network. Yet, instead of projecting the mean and variance in kin-number, we take inspiration from the methods of Tuljapurkar et al. (2020) and seek a discrete probability distribution to reflect kin-number. In more detail, we would like to derive the probabilities that Focal experiences a certain integer number of kin, by age of kin and age of Focal. For example in relation to Figure 1, consider Focal’s older sisters (i.e., **m**). Suppose that Focal is aged *y*. With a set of probabilities, Focal will experience exactly zero, one, two, or more older sisters who are of exact age *s* (where obviously *s > y*).

**Figure 1:**
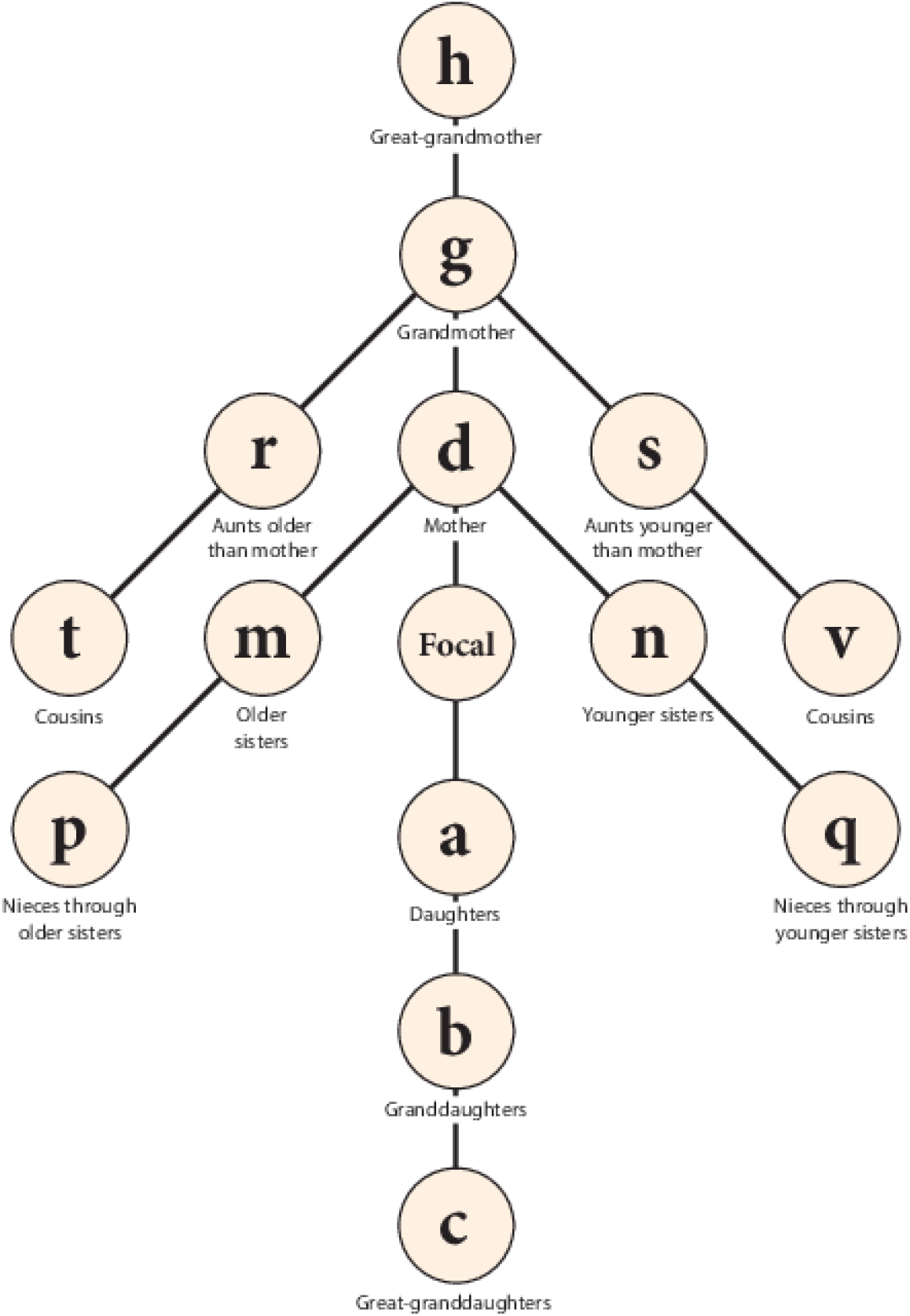
The diagram and notation of kin used in the formal models of Caswell (see e.g. Caswell, 2019, passim).

For any generic of kin pictured in Figure 1, we write the probability mass function (pmf),

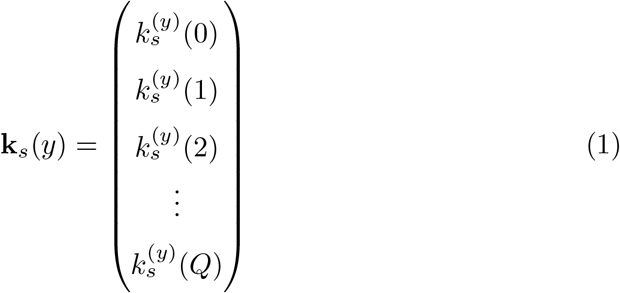

to represent the number distribution of Focal’s kin of type *k* who are of age *s* when Focal is of age *y*. The entries of Eq (1) give the probabilities that Focal experiences some number of that kin: the first, 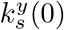 provides the probability that Focal does not have any of the particular kin, the second, 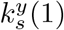 provides the probability that Focal has exactly one of such kin, and the third, 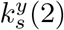,provides the probability that Focal has exactly two of this kin. We must impose a biologically reasonable upper bound *Q*, which acts to limit the maximum lifetime number of the particular kin. As illustrated through Eq (1), the last, or (*Q* + 1)-th entry 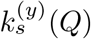, gives the probability that Focal has *Q* kin of type *k*, who are of age *s* when Focal is aged *y*.

### 1.2 Algebraic notation

The framework presented in this article includes mathematical derivations, some of which may be non-obvious. Here we define the main algebraic objects of interest and summarise the notation used throughout this paper. Thus, stochastic matrices are denoted using blackboard bold –𝔸, other matrices are denoted boldface upper-case **A**, vectors lower-case boldface **a**. Discrete probability distribution functions are denoted by boldface Greek symbols, e.g., ***ψ***. We denote the unit vector with *i*-th entry one by **e**_*i*_. When possible matrix entries will be given by lower-case letters, e.g., *a*_*i,j*_ the *i, j* entry of **A**, but when notation is a pain – for instance when the matrix is a function of parameters, **A**(*x*) – we may use [**A**(*x*)]_*i,j*_. The transpose of a matrix is denoted by †.

A discrete convolution of two functions *f* and *g* defined on the integers is given by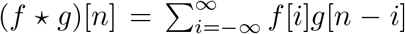. For two distributions ***ψ***_1_ ∈ ℤ ^n^ and ***ψ***_2_ ∈ ℤ ^n^ we write the discrete convolution as ***ψ***_1_ ⋆ ***ψ***_2_ with *m*-th entry defined through (*ψ*_1_ ⋆ *ψ*_2_)[*m*] = ∑ _i_ *ψ*_1_(*i*)*ψ*_2_(*m* − *i*). Over a set of independent distributions 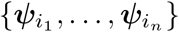 we write

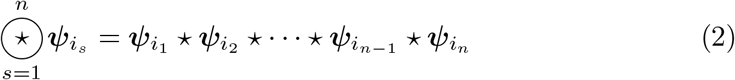

and to represent the *n*-th convolution power of a distribution, we write

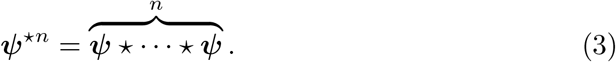

We provide a refresher (with an application to kinship) on the operation of discrete convolution in Appendix C. Lastly, we denote function composition by (*f* º *g*)(*x*) = *f*(*g*(*x*)), and moreover, *f*^[*n*]^(*x*) = (*f* º · · · º *f*)(*x*) as the *n*-times composition of *f*. The ordered composition of functions {*f*_*i*_, *i* = 1, …, *n*} is denoted by

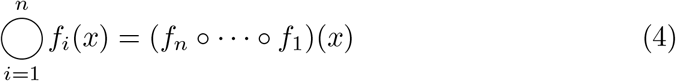

## 2 Model ingredients and assumptions

Here, we outline our assumptions and formally construct a model. Recall that we write **k**_*s*_(*y*) to represent a (*Q* + 1)-dimensional kin-number pmf with (*j* + 1)-th entry giving the probability that Focal has exactly *j* = 0, 1, …, *Q* kin of age *s* when she is age *y*. For consistency henceforth, we refer to the random variable *j* as “kin-number-class”.

### 2.1 Demographic dynamics

Consider a female population structured by age-classes *i* = 1, 2, …, *n*. Within each age-class *i*, the number of births to an individual is a random variable *F*_*i*_ ∈ {0, 1, …} with probabilities {*ψ*_*i*_(0), *ψ*_*i*_(1), …}. Define the vector ***ψ***_*i*_ = (*ψ*_*i*_(0), *ψ*_*i*_(1), …)^*†*^. Within each age-class *i*, individuals’ survival to age *i* + 1 are defined through Bernoulli random variables, *U*_*i*_ ∈ {1, 0} (where 1 represents survival and 0 death) with respective probabilities {*u*_*i*_, 1 − *u*_*i*_}.

Using *f*_*i*_ = 𝔼 [*F*_*i*_] = _j_ *jψ*_*i*_(*j*) and *u*_*i*_ = 𝔼 [*U*_*i*_], we obtain the standard Leslie matrix (Allen, 2010):

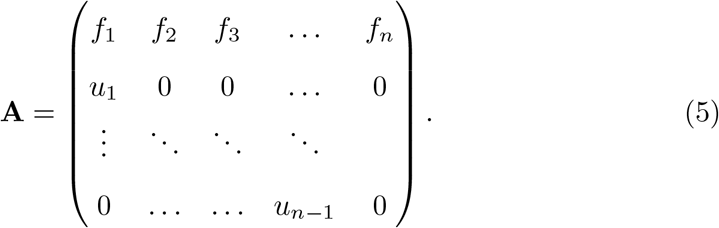

A population vector **x** = (*x*_1_, *x*_2_, …, *x*_*n*_)^*†*^ with *x*_*i*_ representing the number of individuals in age-class *i* is projected from one time to the next through **x**_*t*+1_ = **Ax**_*t*_. Given some initial population structure, **x**_0_, we see **x**_*t*_ = **A**^*t*^**x**_0_ with large-time behaviour (*t >>* 0) resulting in demographic stability: the relative sizes of each age-class become proportional to to the stable age distribution, **w** = (*w*_1_, *w*_2_, …, *w*_*n*_)^*†*^, where **Aw** = *λ***w** (with ||**w**||_1_ = 1). Here, *λ* is the population growth rate, and the stable reproductive values in the population are given by **v** where **A**^*†*^**v** = *λ***v** (we appropriately normalise ||**v**^*†*^**w**||_1_ = 1).

### 2.2 Defining kin of Focal: her *q*-th ancestor and their *g*-th descendant

Following framework of Pullum (1982) we define Focal and kin through a common ancestor. Let *q* be the number of generations that separate Focal from her ancestor. Let *g* be the number of generations that separate Focal’s kin from the ancestor. Define *b*_*i*_ as the age at which Focal’s (direct) *i*-th generation ancestor gives birth to Focal’s (*i* − 1)-th generation ancestor (e.g., *b*_2_ is the age at which Focal’s grandmother gave birth to Focal’s mother), and 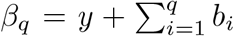 to represent the age of Focal’s *q*-th ancestor when Focal is aged *y*. Note that *β*_*q*_ may be biologically unrealistic but this is not of importance within our framework. As we see below, we use an ancestor’s present-time age as a means to establish their age when reproducing a collateral kin of Focal.

We define *s*_*i*_ as the current age of Focal’s kin, related to Focal as the *i*-th generation descendant of Focal’s *q*-th ancestor. For instance if *q* = 2, then *s*_2_ is age of Focal’s cousin when Focal is age *y*. Note that *s*_*i*_ may be biologically unrealistic. These quantities are used to derive the ages at which descendants of Focal’s *q*-th ancestor reproduce. For instance, Δ_*i*_ = *s*_*i*−1_ − *s*_*i*_ gives the age at which the (*i* − 1)-th generation descendant produces the *i*-th generation descendant. As mentioned above, we also use *s*_*i*−1_ to derive the ages of ancestral reproduction. For instance, *β*_*q*_ − *s*_1_ gives the age at which Focal’s *q*-th ancestor produced a first-generation descendant that is not a direct ancestor of Focal.

We denote the probability distribution for Focal’s [*g, q*] kin which are of age *s*_*g*_ when Focal is age *y* by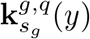, and term this an “age-specific” pmf. We denote the kin-number probability distribution, over all possible ages of kin when Focal is *y*, by 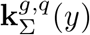. We refer to this as a “total” kin-number pmf.

### 2.3 The probable ages of reproduction

Here we probabilistically predict the age of Focal’s *q*-th ancestor when she gives birth to Focal’s (*q* − 1)-th ancestor. Let *ρ*_*x*_ be the probability that if we randomly sample a newborn from the population, her mother is of age *x*. In a stable demography, the stable population age-structure is proportional to *λ*^−*i*^*u*_1_ … *u*_*i*−1_ (Goldman, 1978). Hence, *ρ*_*x*_ = *u*_1_ … *u*_*x*−1_*f*_*x*_*λ*^−*x*^*/𝒩* where 𝒩 is a normalization constant, which from the Euler-Lotka equation: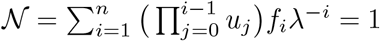.

As such, we see *w*_1_*u*_1_ … *u*_*x*−1_ = *λ*^*x*−1^*w*_*x*_ and thus *ρ*_*x*_ = *f*_*x*_*w*_*x*_*/*(*λw*_1_). Note that *ρ*_*x*_ recovers the (1, *x*)−th entry of the transition matrix of the so-called genealogical Markov chain associated with the population (Demetrius, 1975; Tuljapurkar, 1993). In a stable demography the conditional probability that Focal’s *q*-th ancestor was age *b*_*q*_ when giving birth to Focal’s (*q* − 1)-th ancestor (conditional on other ages of ancestor reproduction *b*_*i*_, *i* = 1 …, *q* − 1) is 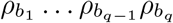 and the unconditional probability is 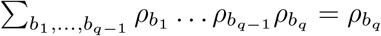

### 2.4 How mortality effects the probabilities of experiencing *j* kin: The matrix U

Here we seek to project a pmf describing the number distribution for a given kin of age *s*^*′*^, conditional on survival, to a pmf describing the number distribution of these kin when they are aged *s* (for *s*^*′*^ *< s*). To do so, we construct a matrix 𝕌 (*s*^*′*^, *s*) ∈ ℝ ^(Q+1)*×*(Q+1)^. Independent of Focal’s age, the role of this matrix is to act on the probabilities 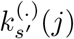 that Focal has *j* = 0, 1, …, *Q* kin of age *s*^*′*^, and to yield probabilities that Focal has 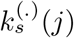 kin of age *s*. That is the action of the matrix is to produce a pmf describing kin-numbers, conditional on survival from age *s*^*′*^ to age *s*.

We formally construct the matrix 𝕌 (*s*^*′*^, *s*) as follows. First, note that each kin independently experiences the same probability 1 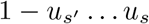 of death between age *s*^*′*^ and age *s*. Introducing the probability 𝒰 (*j, l, s*^*′*^, *s*) that out of *j* kin, some *l* survive from age *s*^*′*^ to age *s*:

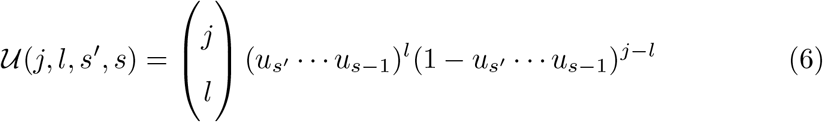

the (*i, j*) entries of 𝕌 are defined through

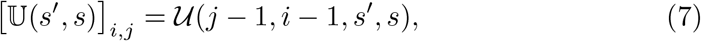

and represent the probabilities that out of *j* − 1 kin, *i* − 1 survive from age *s*^*′*^ to age *s*. The (1, 1) entry of 𝕌 (*s*^*′*^, *s*) is 1, and acts on the *j* = 0 kin-number-class. This ensures that the probability that there is no kin of age *s*^*′*^ contributes a one-to-one correspondence to the probability that there is no kin of age *s*. The diagonal of 𝕌 (*s*^*′*^, *s*) represents the probabilities that all kin survive, while the top-row (excluding the (1, 1) entry) define the probabilities that all of the *j* = 1, 2, …, *Q* kin die (resulting in Focal having zero of such kin of age *s*).

To summarise: the matrix 𝕌 projects kin-number pmfs of age *s*^*′*^ to age *s*. The matrix is both upper-triangular (since survival cannot act to increase kin number) and column-stochastic (see Appendix B). The latter property distributes the total probability of death from any one kin-number-class *j*^*′*^, into kin-number-classes 0 ≤ *j*^*′′*^ *< j*^*′*^, while maps the probability of survival into itself. We adopt the convention that 𝕌 (*s*_1_, *s*_2_) = **I**, ∀*s*_2_ *< s*_1_.

### 2.5 The probabilities that out of *j* kin, *l* offspring are produced: The matrix F

Here we seek to find pmfs which describe the number of newborns created through the age-specific reproduction of one of Focal’s kin. While we are dealing with the reproduction of a direct ancestor of Focal, *j* = 1 (Focal can only have one of these kin). Assume that this ancestor is of age *b*_*i*_. Then we know that its reproduction pmf is simply given through 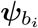.However, if we are dealing with the reproduction of kin who are defined through a pmf with kin-numbers *j* = 0, 1 …, *Q* (e.g., sisters), we have to create a matrix function 𝔽 (*s*) (a function of the kin’s age *s*) which projects the pmf of kin onto a pmf of their offspring.

Because reproduction is independent, the probability that *j* kin produce *l* offspring is the *l*-th entry of the *j*-th power convolution of *ψ*_*s*_ (see Appendix C):

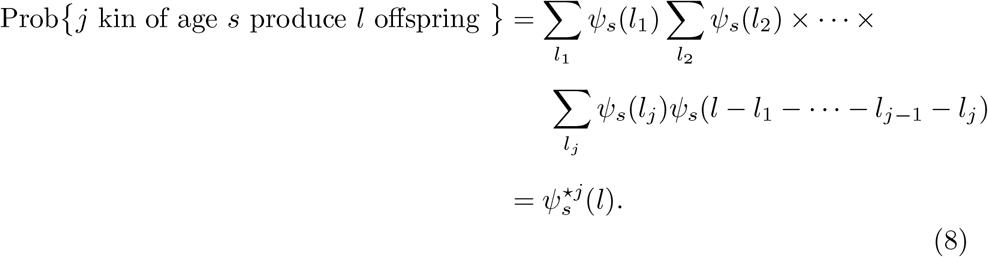

Using this information, one can create a column-stochastic matrix which projects the kin-number distribution of producer kin of age *s* onto a kin-number distribution of offspring kin:

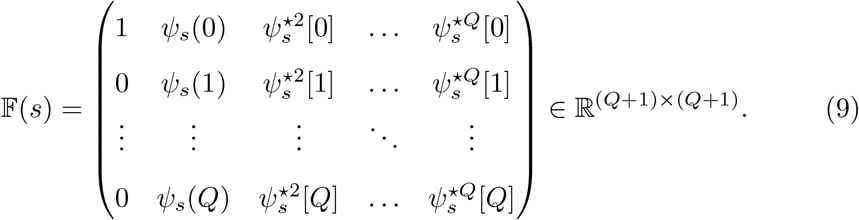

Through 𝔽 (*s*)**k**_*s*_ we produce a pmf of offspring number encapsulating the reproduction of an arbitrary kin-type **k**_*s*_ of age *s*. The first entry of 𝔽 (*s*)**k**_*s*_ reads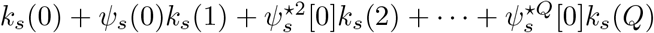, which one interprets as the probability that there are no producer kin (who do not reproduce), plus the probability that there is one kin who does not have offspring, plus the probability that there are two kin and neither have offspring, …, plus the probability that there are *Q* kin and none of them have offspring. The second entry is 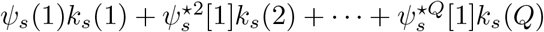 which reads the probability that there is one kin who has one offspring, plus the probability that there are two kin and between them produce one offspring (irrespective of ordering), …, plus the probability that there are *Q* kin and between them all one offspring is produced. The last entry is 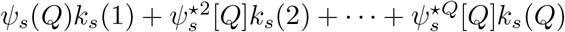 and reads the probability that there is one kin who has *Q* offspring, plus the probability that there are two kin and between them *Q* offspring are produced, …, plus the probability that there are *Q* kin and between them all *Q* offspring are produced.

### 2.6 Total kin versus age-distributed kin

Recall from Section 2.2, that **k**_*s*_(*y*) represents an age-specific kin-number pmf for Focal’s kin of age *s* when Focal is of age *y*, whereas **k**_∑_(*y*) yields a pmf for total kin (over all their possible ages) when she is *y* years old. The 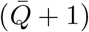-th entry of the latter distribution gives the full probability that Focal has 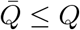 kin. If Focal can have any integer number of kin at any point in her life (i.e., a non-ancestor) then the following theorem (a proof of which is given in Appendix C) provides the pmf for **k**_∑_(*y*):

#### Theorem 1.

*For kin of Focal which are not direct ancestors, the probability that Focal of age y has exactly* 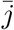 *kin whose ages can range between z*_1_ *and z*_*n*_ *is given by the* 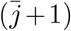*-th entry of* 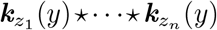.*Moreover*,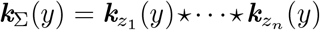.

As a special case, if Focal can have at most one kin at any point in her life (i.e., an ancestor) then we have the following lemma which describes the pmf for **k**_∑_(*y*):

#### Lemma 1.

*For kin of Focal which are direct ancestors, the probability that when Focal is aged y this ancestor is alive is given by p* =∑ _s_[***k***_*s*_(*y*)]_2_ *(where* [.]_*j*_ *represents the j-th entry), resulting in* ***k***_∑_(*y*) = (1 − *p, p*, 0, …, 0)^*†*^.

Lemma (1) is intuitive since Focal has, with probability 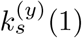 (the second entry of **k**_*s*_(*y*)), one living ancestor of age *s*. The total probability that the ancestor is alive is given by the sum over all biologically feasible ages *s* that the ancestor may be.

## 3 Towards the kin-formulae

Using the above ingredients and observations, we now provide the explicit formulae for kin. We break down the formula into four theoretically distinct regimes. These regimes correspond to the branches of the tree in Figure 1. First, branches which extend right (older lineages) reflect kin which descend through older sisters of Focal’s (*q* − 1)-th ancestor. Second, branches which extend left (younger lineages) reflect kin which descend through younger sisters of Focal’s (*q* − 1)-th ancestor. Third, branches extending down reflect Focal’s descendants. Lastly, branches extending up reflect direct ancestors of Focal. In the next sections we respectively deal with these cases in turn, while the particular case whereby kin descend through same-age-class sisters of Focal’s (*q* − 1)-th ancestor is dealt with in Appendix D. A more detailed break down of the formulae and how they relate to the established kinship model of Caswell (2024) is given in Appendix E. We provide an illustrative application of our framework in Section 4.

In order to simplify the exposition moving forwards, we introduce the operator,

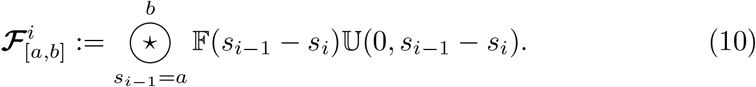

which acts on the probability mass function of a kin-number distribution of newborn kin for an arbitrary (*i* − 1)-th generation descendant of Focal’s *q*-th ancestor. The operator, conditional on survival, takes the convolution over the reproduction of these kin between the age *s*_*i*−1_ − *s*_*i*_ = *a* and *s*_*i*−1_ − *s*_*i*_ = *b*. We set *n* as the minimum fertile age and 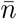 the maximum.

### 3.1 Kin which descend through older sisters of Focal’s (*q* − 1)-th ancestor

Focal’s [*g, q*] kin which descend from an older sister of Focal’s (*q* − 1)-th ancestor, and which are of age *s*_*g*_ when Focal is age *y* are denoted by 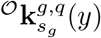.Introducing the reproduction pmf of Focal’s *q*-th ancestor at birth of their first descendent, older than Focal’s (*q* − 1)-th ancestor,

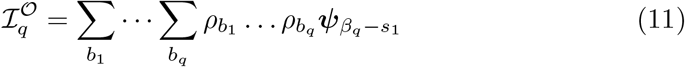

and applying the operator defined in 10 to

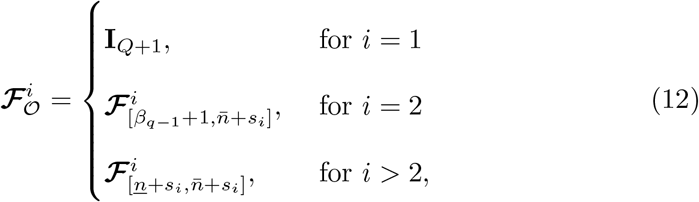

the age-number probability distribution of these kin can be written as a composition of operators:

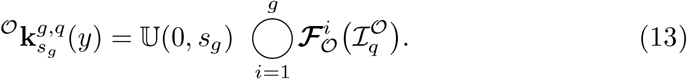

In Eq (13), if we consider *g* = 1 then 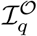 recovers a discrete probability distribution for the number of newborn kin, born to Focal’s *q*-th ancestor *before* the birth of Focal’s (*q* − 1)-th ancestor. Focal’s *q*-th ancestor produced Focal’s (*q* − 1)-th ancestor at age *b*_*q*_; at age *β*_*q*_ − *s*_1_ she had *j* = 0, 1, …, *Q* many newborns with probabilities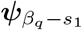. Summing over *b*_*i*_, *i* = 1, …, *q* yields the complete probability distribution of these newborns. Pre-multiplication the pmf of newborns through 𝕌 (0, *s*_1_) procures a pmf of these kin at age *s*_1_ (when Focal is age *y*).

Otherwise, if *g >* 1 then the limits of *s*_1_ (i.e. when *i* = 2) over which the convolution of reproductive ages are taken, defined through Eq (12), ensure that kin descend through older sisters of Focal’s (*q* − 1)-th ancestor. For *i >* 2 convolutions over reproductive ages have no constraints.

For further details we refer the reader to the most simple examples of Eq (13), described with annotations in Appendix E.1 and Appendix E.2.

### 3.2 Kin which descend through younger sisters of Focal’s (*q* − 1)-th ancestor

We denote by 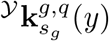, Focal’s [*g, q*] kin which descend from a younger sister of Focal’s (*q* − 1)-th ancestor, and which are of age *s*_*g*_ when Focal is aged *y*. Introducing the reproduction pmf of Focal’s *q*-th ancestor at birth of their first descendent, younger than Focal’s (*q* − 1)-th ancestor

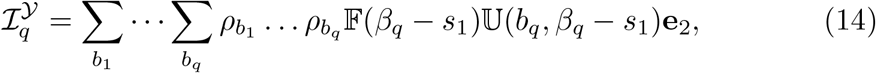

and again applying the operator defined in 10,

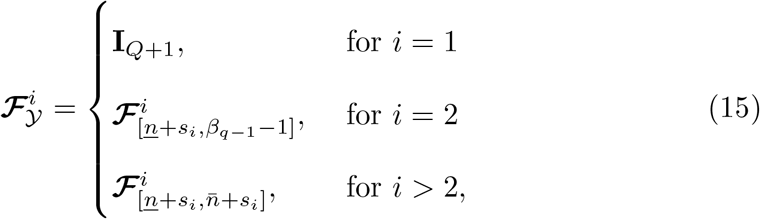

the age-number probability distribution of these kin is given by the composition:

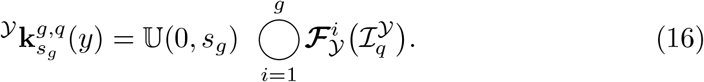

In Eq (16), first note that we know with certainty that Focal’s *q*-th ancestor produces Focal’s (*q* − 1)-th ancestor at age *b*_*q*_. Here the ancestor has a pmf defined through a unit vector with mass in the 1 kin-number-class, **e**_2_. If *g* = 1 then we recover 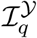 as defined in Eq (14). The summand term produces, through pre-multiplication by 𝕌 (*b*_*q*_, *β*_*q*_ − *s*_1_) a distribution representing the probabilities that Focal’s *q*-th ancestor was alive at the age when she produced Focal’s kin *β*_*q*_ − *s*_1_, and through pre-multiplication by 𝔽 (*β*_*q*_ − *s*_1_), a probability distribution representing their offspring, born *after* the birth of Focal’s (*q* − 1)-th ancestor. Summing over all probable *b*_*i*_ yields a complete pmf of newborns. Pre-multiplication through 𝕌 (0, *s*_1_) then calculates the probabilities than between birth and age *s*_1_ (when Focal is aged *y*) the kin survive.

While *g >* 1, the case of *i* = 2 is diametrically opposed to Section 3.1; the limits of *s*_1_ over which the convolution of newborn-number probability distributions are taken, are constrained such that *s*_1_ − *s*_2_ ≤ *β*_*q*−1_ − 1 − *s*_2_. This ensures that kin descend through younger sisters of Focal’s (*q* − 1)-th ancestor. For *i >* 2 convolutions over reproductive ages have no constraints.

The most basic applications of Eq (16) derive Focal’s younger sisters, and then nieces through these younger sisters. These cases are respectively described with annotations in Appendix E.5 and Appendix E.6.

### 3.3 Descendants of Focal

Descendants of Focal can be calculated through Eq (13) with *q*:= 0. Recall *s*_*i*_ is the age of Focal’s *i*−th direct descendant. By appealing to the operator in 10we define

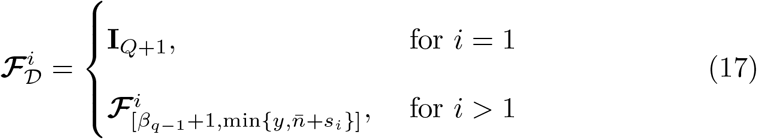

and find the pmf of Focal’s *g*−th descendant of age *s*_*g*_ when Focal is *y*, as

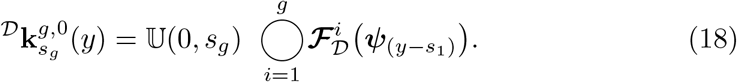

Eq (18) tells us that Focal (who is assumed immortal) will have *g*-generation descendants of age *s*_*g*_ if she gives birth to offspring at age *y* − *s*_1_ who in turn survive to *s*_1_ − *s*_2_ and reproduce at this age, continuing in this manner up to Focal’s (*g* − 1)-th generation descendant, who survives from birth to age *s*_*g*−1_ − *s*_*g*_ at which point producing Focal’s *g*-th descendant (who survives to age *s*_*g*_). Because at each generation *i* of descent, Focal can have multiple descendants, the (*i* + 1)-th generation descendants are given by the convolution of pmfs of the *i*-th generation over the fertile age range (recall Theorem (1)).

### 3.4 Ancestors of Focal

Descendants of Focal correspond to the case whereby *g*:= 0. Recall *q*_*i*_ is the age of Focal’s *i*-th direct ancestor when producing the (*i* − 1)-th. We obtain the closed form equation for the pmf of Focal’s *q*-th ancestor of age *s*_0_ when Focal is *y*, through

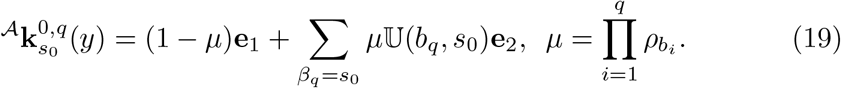

The sum is taken over all possible arrangements of the ages *b*_*i*_ at which Focal’s *i*-th ancestor produced Focal’s (*i*−1)-th, such that the age of Focal’s *q*-th ancestor when Focal is *y, β*_*q*_ = *b*_1_ + … *b*_*q*_ + *y* is equal to *s*_0_. The vector *µ***e**_2_ in Eq (19) assign binomial probabilities that Focal’s *q*-th ancestor is indeed defined through the genealogical sequence of *b*_*i*_. We project the *q*-th ancestor to survive from age *b*_*q*_ – when last we knew it was in existence (producing (*q* − 1)-th ancestor) to survival to her age *s*_0_ = *b*_1_ + *b*_2_ + · · · + *b*_*q*_ + *y* when Focal is *y*. The vector (1 − *µ*)**e**_1_ re-normalises by adding to the pmf, the probability that Focal’s *q*-th ancestor was not defined by the sequence of *b*_*i*_ (i.e., is not of exact age *s*_0_).

## 4 Application

Here, the model is applied. We source single year of age, period based (from 1974), UK fertility and mortality data from the Human Fertility and Mortality Databases (Human Fertility Collection, 2024; Human Mortality Database, 2024). We make the simplifying assumption that births follow a Poisson distribution. Thus, with an age-specific fertility rate *f*_*i*_, the distribution ***ψ***_*i*_ has entries giving the probabilities of having *k* offspring: 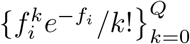. Our assumption here in no way restricts the model, but is simply used for progress. We could equally use, for example, empirical distributions of multiple births for single year of age data. Deaths follow a Bernoulli distribution: we write {*u*_*i*_, 1 − *u*_*i*_} for the probabilities of survival from age *i* to *i* + 1 (or not).

We set *Q* = 6 as an upper bound for the maximum lifetime kin-number. Of course we could use a larger upper limit, however, it would be unlikely for an individual to experience more than 6 sisters, aunts or cousins, during their life^1^. In Appendix A we also explore the validity of the below results by comparing them to a micro-simulation. All code and data used, as well as a comparison of the mean field results of this model to (Caswell, 2019), can be found at [link redacted for anonymous review]

### 4.1 Unconditional distributions of kin

#### 4.1.1 Sisters and cousins

Using Eq (13) and Eq (16) respectively, we are able to produce age-specific kin-number pmfs for Focal’s younger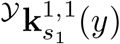, and older 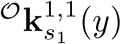, sisters of age *s*_1_, for each age *y* of Focal’s life. For each feasible age of Focal’s kin, *s*, these provide probabilities that Focal has exactly *j* = 0, 1, …, *Q* of kin that age. Given these distributions, it is a simple case of applying Theorem (1) to obtain the overall distributions of kin (of all possible kin ages when Focal is age *y*):

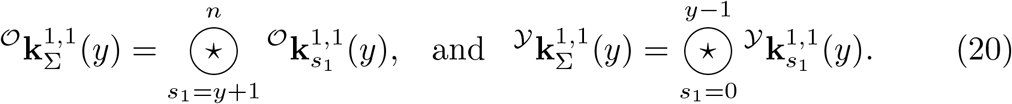

The complete pmf for the kin-number distribution of sisters Focal has is given by the convolution of the distributions in Eq (20). Using *g* = *q* = 2, we mathematically decompose Focal’s cousins as descending from younger sisters of Focal’s mother 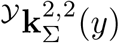,and older sisters of Focal’s mother 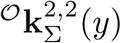.The complete kin-number pmf of cousins is given by the convolution of these distributions.

Figure 2 illustrates the age-specific pmfs describing the total numbers of sisters and cousins of Focal, for two ages in Focal’s life-course; when she is aged 20 and aged 50. Plotted are expected kin-numbers, the standard deviation in kin-number, and the skewness in kin-number. Because we consider probabilities that Focal experiences some number of kin of a specific age, nearly all the mass appears in the first (i.e., 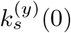 entries of the pmfs: the distributions display a heavy (positive) skew.

**Figure 2:**
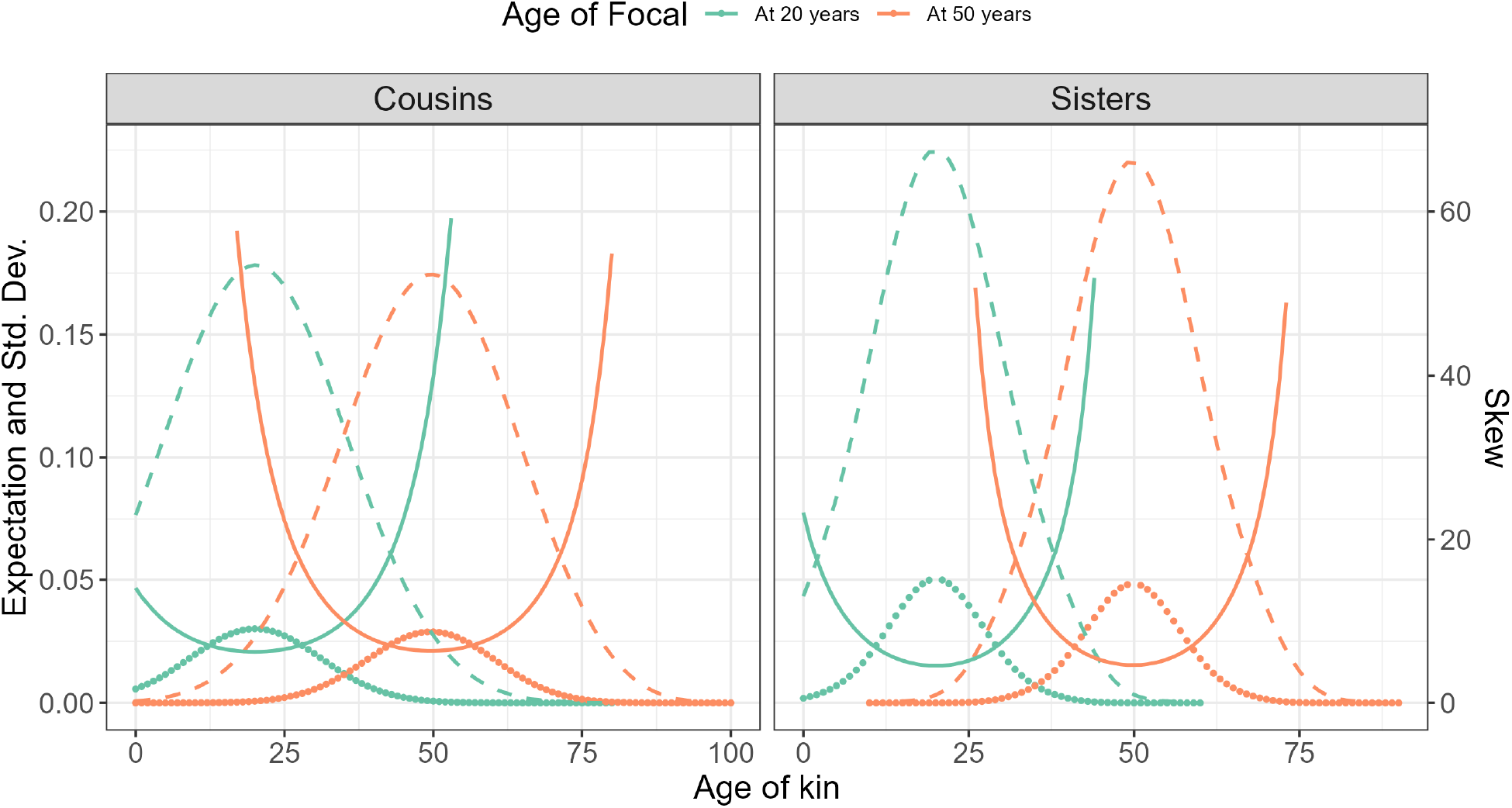
Theoretically predicted age-specific pmfs for Focal’s sisters (right) and cousins (left) when Focal is aged 20 (turquoise) and 50 (orange). Points show expectations, dashed lines show standard deviation, and solid lines show skew (separate *y*-axis). For ease of visualisation, we omit values ≥60 when plotting the skew.

Figure 3 illustrates our theoretically predicted pmfs for the total number sisters and cousins of Focal when she is aged 20 and 50. We predict that when Focal is aged 20 she has no sisters with a probability 0.42 and no cousins with a probability 0.46. When Focal is aged 50, we predict that she has no sisters with a probability 0.43 and no cousins with a probability 0.50. Also presented are statistics derived from the first, second, and third moments of the kin-number distributions.

**Figure 3:**
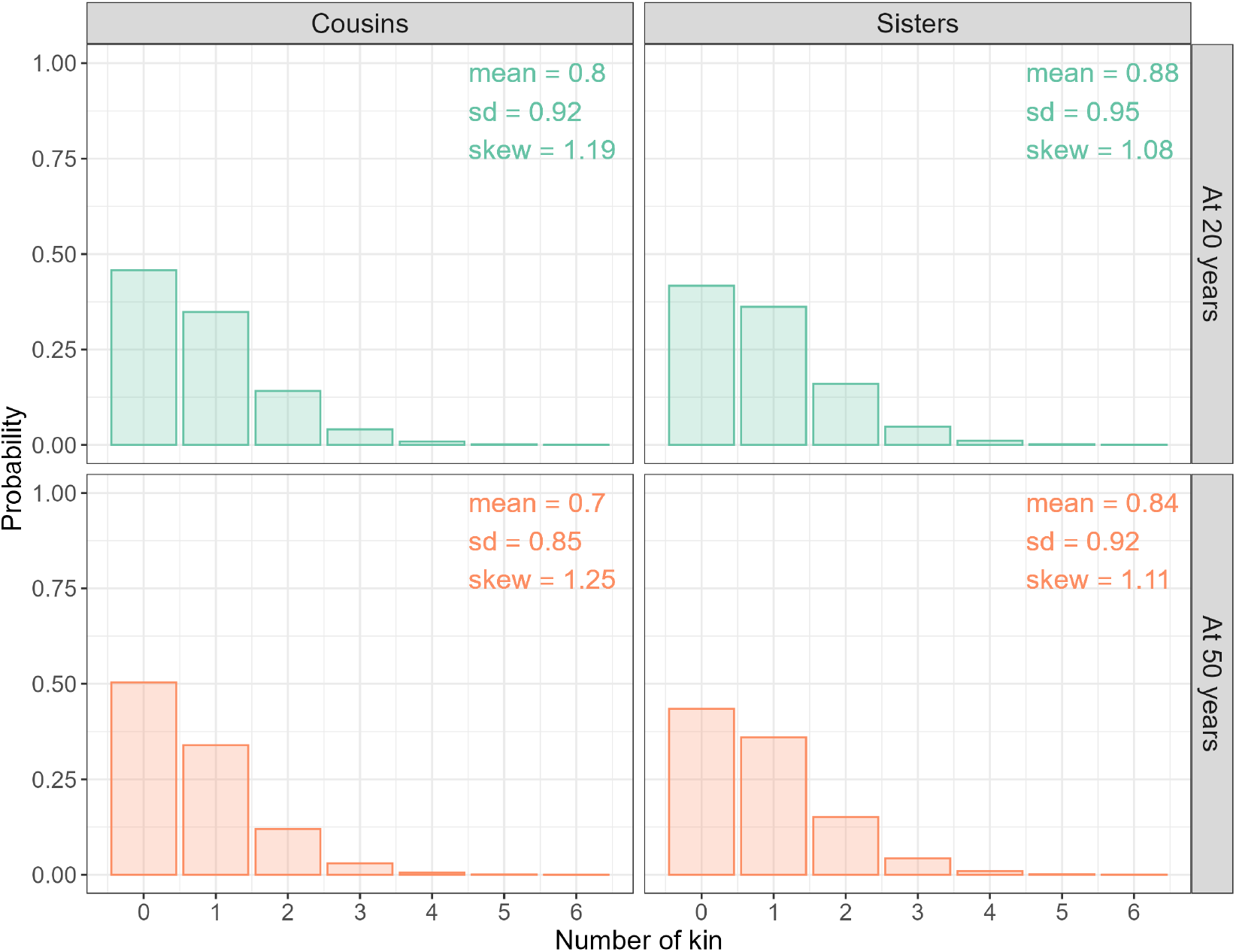
Accumulated-kin pmfs for Focal’s cousins (left) and sisters (right) when Focal is aged 20 (top row) and 50 (bottom row). Bars show the probability that Focal has a given number of kin.

In Appendix A, we compare the above predictions for the accumulated number-probability distributions of kin of Focal to a direct stochastic simulation (see Figure A-1 and Figure A-2).

#### 4.1.3 Ancestors and descendants

Applying Eq (19) we obtain 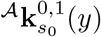, the age-specific pmf for Focal’s mother of age *s*_0_ when Focal is *y*. Through Eq (18) we obtain 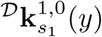,the age-specific pmf for Focal’s daughters of age *s*_1_ when Focal is age *y*. These distributions are visualised in Figure 4 where we plot the expected kin-numbers, as well as the standard deviation and skewness in kin-number. We focus on two different ages in Focal’s life course; when she is aged 20 and 50. We again observe that the distributions are heavily skewed.

**Figure 4:**
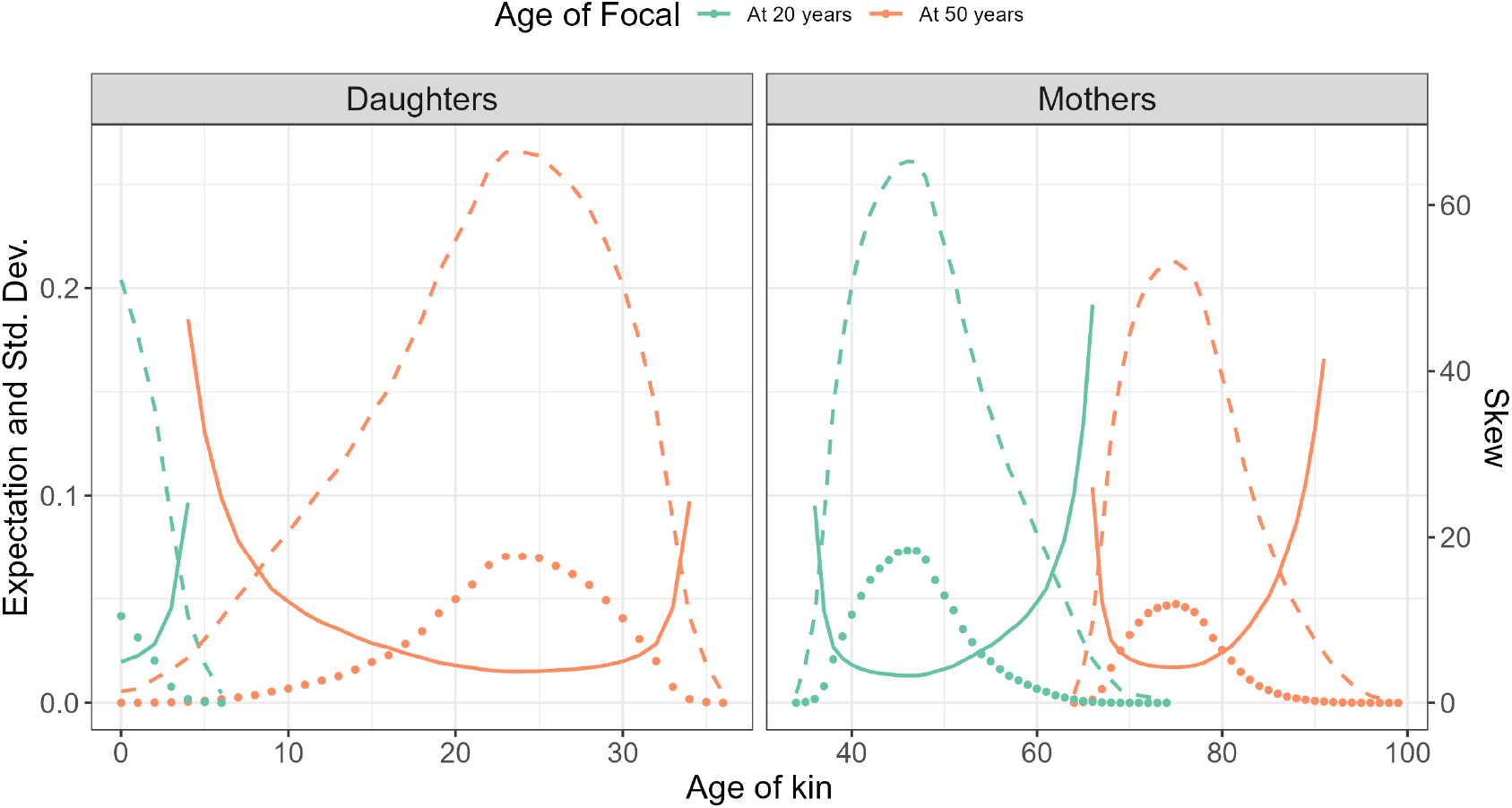
Theoretically predicted age-specific pmfs for Focal’s daughters and mother when Focal is aged 20 (turquoise) and 50 (orange). Left; daughters, right; mother. Points show expectations, dashed lines show standard deviation, and solid lines show skew (separate *y*-axis). For ease of visualisation, we omit values ≥ 50 when plotting the skew.

In the same vein as in Section 4.1.1, we obtain the pmf of total kin for Focal of ages 20 and 50 by applying Theorem (1):

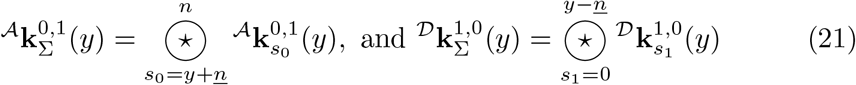

Figure 5 illustrates these pmfs. In each panel, the bars show the probabilities that Focal experiences a number of kin. The left column shows Focal’s daughters, while the right Focal’s mother. Rows show age of Focal. By comparing Focal at ages 20 and 50, we see how the probable numbers of kin change by age of Focal. Regarding daughters, when Focal is aged 20 and early her reproductive cycle, we predict that she will experience one or more daughters with probability 0.098. Contrastingly, at age 50 when Focal has completed fertility, we predict that she will experience one or more daughters with probability 0.606.

**Figure 5:**
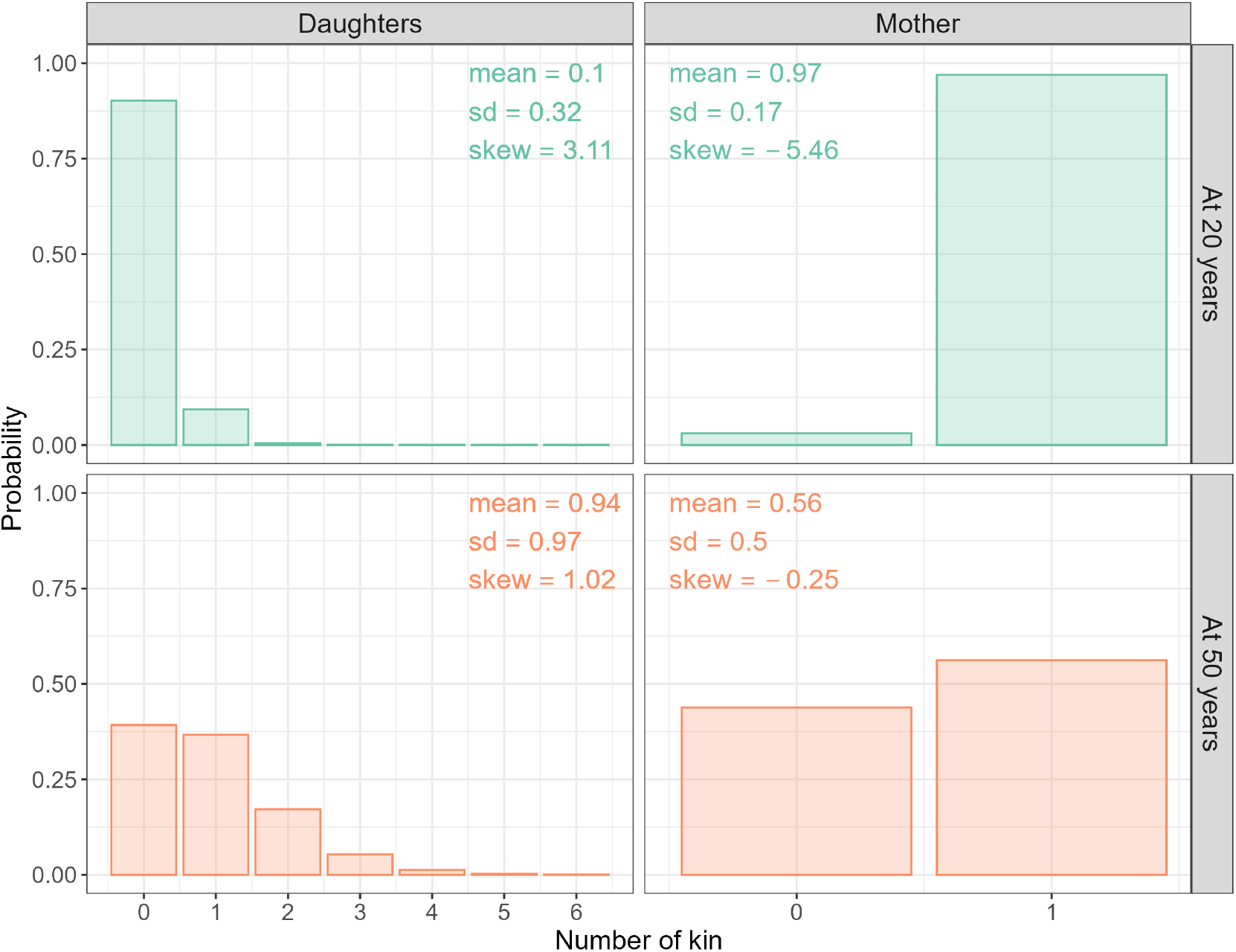
Accumulated-kin pmfs for Focal’s daughters and mother when Focal is aged 20 and 50. Rows show age of Focal. Left; daughters, right; mother. Bars give the probability that Focal has a given number of kin; vertical lines show the expected numbers of kin; horizontal error-bars show expected kin-number ± standard deviation in kin-number.

Also illustrated are the expected numbers of kin: daughters; 0.10 at age 20 and 0.93 at age 50, and mother; 0.97 at age 20 and 0.56 at age 50, the standard deviation in kin-number: daughters; 0.32 at age 20 and 0.97 at age 50, and mother; 0.17 at age 20 and 0.50 at age 50, and the skewness in kin-number: daughters; 3.11 when Focal is aged 20 and 1.02 when Focal is aged 50, and mother; -5.46 when Focal is 20 and -0.25 when Focal is 50.

In Appendix A, we compare our above predictions of how the number of kin Focal changes over her life course to a direct stochastic simulation (see Figure A-3 and Figure A-4)

### 4.2 Conditional distributions of kin

Our method allows one to condition the probability that Focal experiences some number of one kin-type on life-history events pertaining to other kin-types. For example, consider Focal’s sisters. To find the probability distributions for Focal’s sisters, we apply a weighted average of the age-specific reproduction of Focal’s mother, over possible ages when she had Focal. Focal’s mother, however, is one age when having Focal. It is thus of interest to remove the weighting and view the conditional pmfs of Focal’s sisters, assuming certain (error-free) knowledge of Focal’s mother’s age when she had Focal.

#### Illustration: Focal’s sib-ship conditioned on age of mother

Assuming that Focal’s mother was aged *b*_1_ when she had Focal, the respective conditional distributions representing the probability that Focal has *j* = 0, 1, …, *Q* older or younger sisters of age *s* when she is aged *y*, are

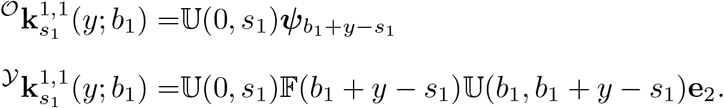

The conditional pmfs of total older and younger sisters (over all possible ages) are

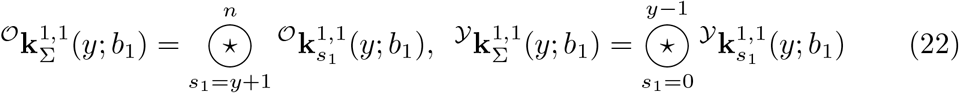

and the convolution of the distributions in Eq (22) gives the complete probabilities that Focal has a sister (“combined” younger or older), conditional on her mother being aged *b*_1_ when she was born. For each 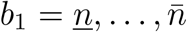,the different probable number of sisters of Focal, as per Eq (22), is shown in Figure 6. The left hand panel shows probabilities that Focal has some number of younger sisters. Here, the younger Focal is, the less time her mother has to produce her younger sisters, and the less likely Focal has such kin. At older ages of Focal, the presence of younger sisters is more likely but depends on whether Focal was born earlier or later in the reproductive ages of the mother. Contrastingly, the middle panel – reflecting probabilities that Focal experiences some number of older sisters – remains invariant with respect to Focal’s age. Here, the probabilities only change with the age at which Focal’s mother had Focal. This is because Focal has accumulated all of her older sisters before birth. The right hand panel gives the probability that Focal has some given number of combined (i.e., younger or older) sisters of any age.

**Figure 6:**
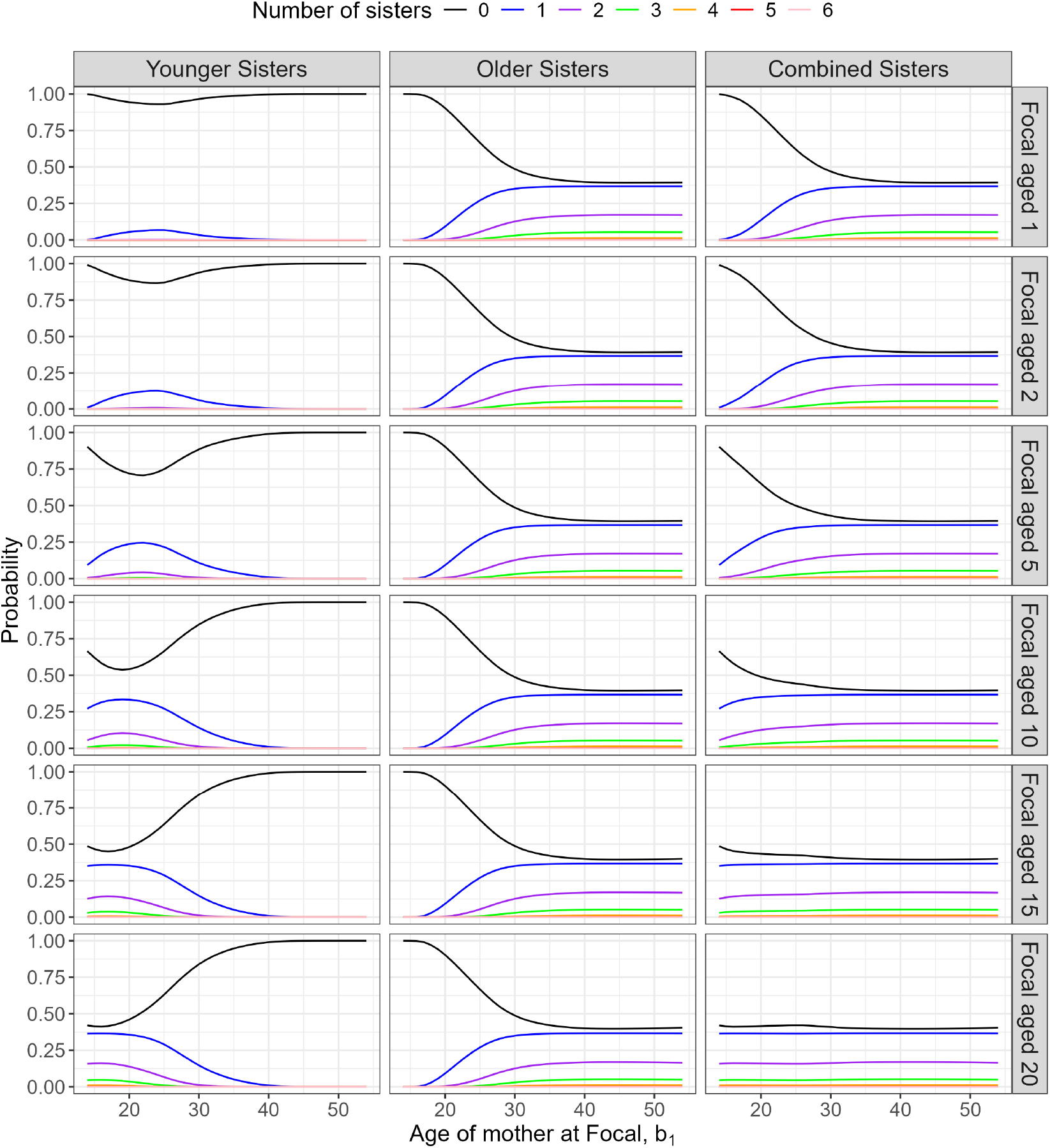
The probable number of sisters of Focal as a function of the age of Focal’s mother, at different times in Focal’s life. Rows give the age of Focal. Left: younger sisters; middle: older sisters; right: combined sisters.

We might also consider the probability that Focal has no sisters. These probabilities once again depend on the age of Focal’s mother when Focal was born; see Figure 7. In the plot, the left panel, illustrating the probability that Focal has no younger sisters, demonstrates that the older Focal’s mother is at birth of Focal, the less likely she experiences such kin. In this case, Focal simply doesn’t accumulate younger sisters since mother is nearing the end of her reproductive interval. The more probable instances of Focal experiencing a younger sister occur when her mother is relatively young when having Focal, and when Focal is currently at an age whereby her mother will have completed reproduction. Interestingly, for Focal up to age 10, if her mother gave birth to her at a very young age (e.g., *b*_1_ = 15) we find that it more probable for Focal to have no younger sisters compared to if Focal’s mother gave birth to her at a mid-fertile age (e.g., *b*_1_ = 25). Because age-specific fertility is unimodal in our example (1974; UK), if Focal’s mother is young when mothering Focal she retains a low probability of reproduction in the immediate years after; thus Focal has to wait before accumulating younger sisters.

**Figure 7:**
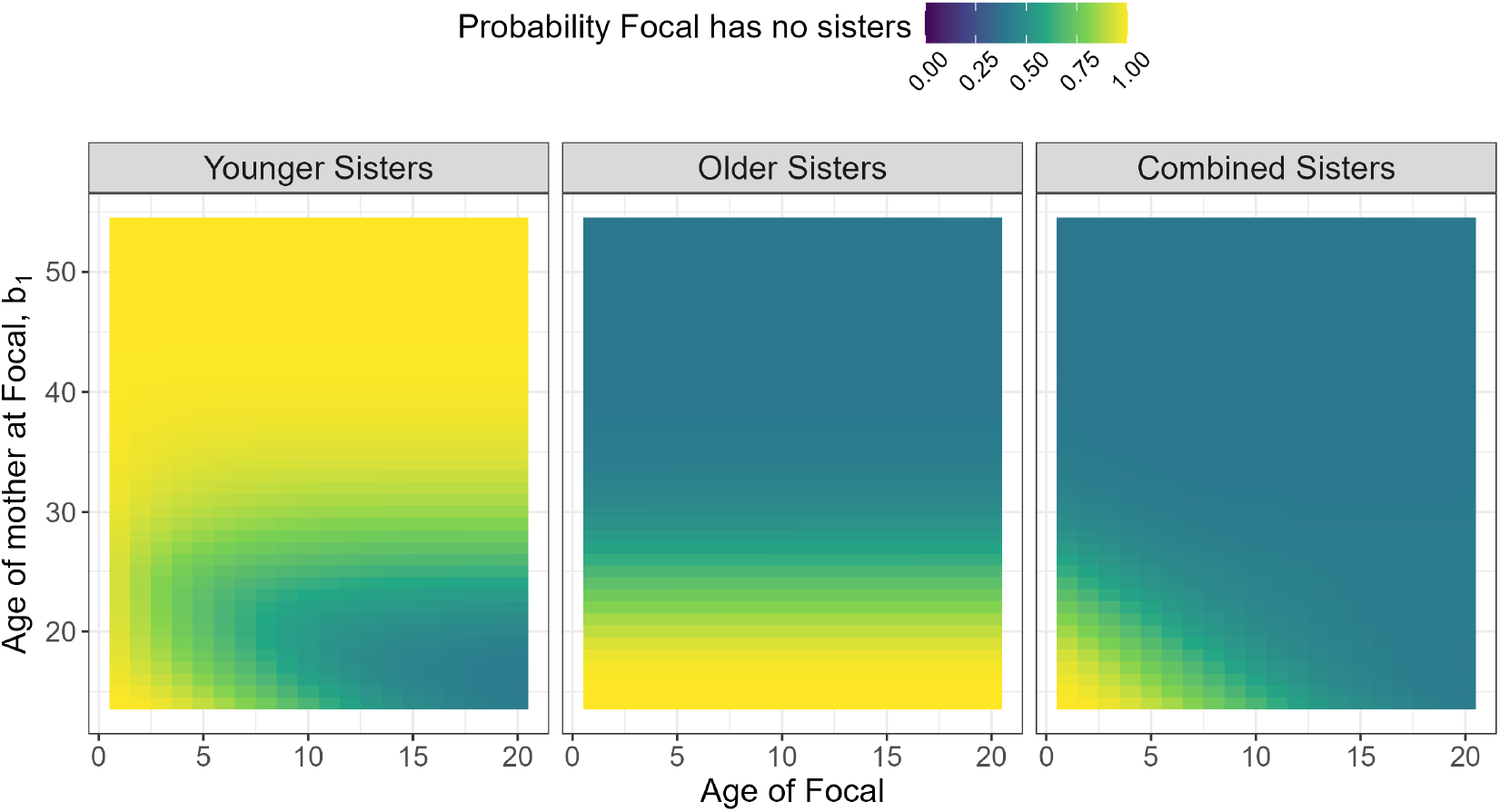
The probability that Focal has no sisters as a function of age, and conditional age of mother at Focal. Left: no younger sisters; middle: no older sisters; right: no younger or older sisters.

Note that in the present example we do not account for birth interval in the fertility modelling, or else we might expect to see more complex patterns in these heat maps, for example indicating that transition to gaining a first younger sister is less likely as Focal ages, separately from the fertility decline as mother ages.

Focal’s sib-ship size *S* (excluding Focal) is a random variable, conditioned on the age of mother at birth of Focal, *b*_1_. Let the variable *X*_*i,j*_ indicate the probability that Focal has *i* older sisters and *j* younger sisters (i.e., *i* + *j* = *S*).

Thus, for each possible sib-ship size, *j* = *S* − *i*, and as such

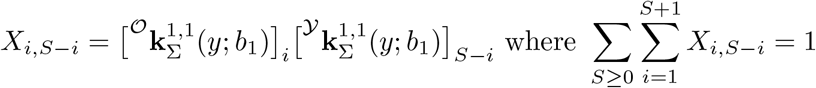

with the right-hand side guaranteed by the conditioning on mother’s age at birth of Focal. The variable *X*_*i,S*−1_ allows us to calculate the probability of Focal’s birth rank, as illustrated in Figure 8. Here we plot birth rank probabilities for Focal of age 20: (i) conditional on mother’s age *b*_1_ (by row) and (ii) as a weighted mix over probable ages of mother (bottom row). Note that each of the rows sum to one because conditional on mother’s age, they provide probabilities of Focal having *S* = 0, 1, …, 6 sisters, and given that, Focal’s birth order. We might also condition on sib-ship size, in which case we would simply re-normalise each panel.

**Figure 8:**
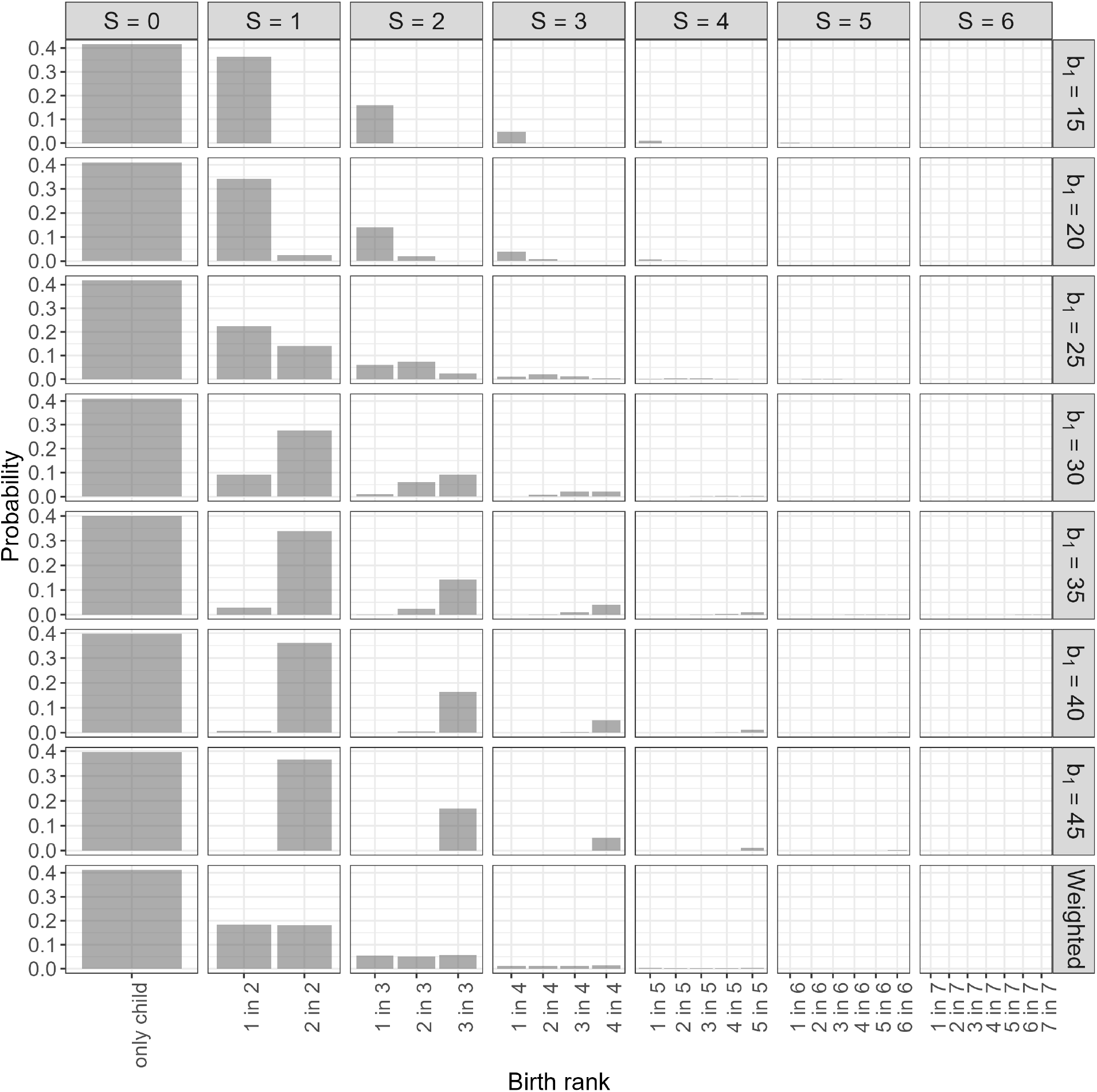
Probability that when 20 years old, Focal experiences *S* sisters, and from them *i* are older; *S* − *i* are younger. Conditioning on the age of Focal’s mother at birth of Focal (*b*_1_) is illustrated by rows (the bottom row shows the weighted average over all *b*_1_). Columns give sizes of sib-ship (*S*).

## 5 Discussion

The research presented in this paper extends and enriches recent advances in modelling demographic stochasticity within kinship networks (Caswell, 2024). Presently, the only published model to acknowledge the above-mentioned stochasticity accounts only for the mean and variance in kin-number. Our proposed frameworks projects a complete probability distribution of the number of kin, from which, higher-order moments or other quantities of interest (such as quantiles) can be readily obtained. Our framework thus provides a more comprehensive analysis of variability in kinship structures. The probabilistic nature of our framework allows conditioning kinship networks on particular life-history events (see Section 4.2). The approach presented here thus allows analytical results regarding demographic uncertainty that approach the richness of those obtained via micro-simulation, providing population distributions of numbers of kin by age of both Focal and kin, for arbitrary kin types.

Our proposed framework treats probability distributions for age-specific mortality and fertility as model inputs. From these, the probability distributions of kin naturally emerge. This innovation allows one flexibility to choose from empirical distributions (e.g., single-year-of-age offspring probabilities) as well as parametric distributions. Contrastingly, fitting empirical distributions using the mean and variance in kin numbers – outputs of the state-of-the-art stochastic kinship model – would be challenging. Another innovation made here is that we reduce the dimensionality of the matrix projections. The size of the state-space here is defined by life-time maximum kin-number, *Q*. Although, as argued below, this does not necessarily offer a computational advance, this feature of our model alleviates the need for big state-space matrices.

In principle, our model utilises the theory of branching processes with incorporated age-structure. We demonstrate the recursive nature of kin-structures, quite similar to the seminal work of Goodman, Keyfitz, and Pullum (1974), however with convolutions of distributions implementing the next generation of kin rather than integrals over scalar expectations. Waugh (1981) provides a way to construct a joint probability generating function (pgf) for generation size over multiple generations in a Galton-Watson process, and then samples a typical “Ego” (rather than Focal) from a fixed generation (see e.g., Section 7 in that paper). In this way, the author elegantly calculates what they refer to as Ego’s “sorority” (sib-ship) size. The addition of age-structure in our framework builds on such work and allows one to calculate probabilities of birth order as well as overall sib-ship size. Moreover, in order to apply the methods proposed by Waugh (1981), one requires an analytic pgf (they use a fractional linear functional form). For progress here, we avoid such restrictions and maintain a discrete probability distribution which is easier to extract from empirical data. From a computational perspective, we cannot compete with the leading matrix-projection model Caswell (2024). The manner in which distributions of collateral relatives are calculated requires summations over ancestral reproductions and convolutions over their descendants. However, our equations are substantially faster to implement than an individual-based micro-simulation model, and in the present context, are able to yield results of comparable complexity. Note that in higher-dimensional settings, for instance whereby additional stages reflect characteristics of kin, incorporating additional complexity into a micro-simulation is likely to be easier than extending our analytical expressions. When comparing the results presented in Section 4 to a direct stochastic simulation (available to explore at [link redacted for anonymous review]) we find overall consonance in results. For kin which form ancestors and descendants in Focal’s family tree, the kin-number distributions are near exact (see Figure A-3 and Figure A-4). Regarding sisters, we find agreement in the expectation of kin-numbers, but subtle differences in the overall pmfs. This results in the variance in the number of sisters being slightly larger in the simulation (see Figure A-1 and Figure A-2). A possible explanation for this discrepancy is Monte-Carlo error. Nonetheless, we cannot rule out the effects of demographic stochasticity. In finite populations, fluctuations in births and deaths will result in the realised proportions of individuals in each age class deviating from the stable distribution. Consider a population of size *N*(*t*) at time *t* structured by age classes *i* so that 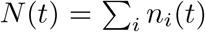.Recall from Section 2.1 that *F*_*i*_ is a random variable representing offspring number for an individual in age-class *i*. Using 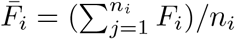 to represent the average age-class reproductive value, the deviation in individual reproduction from the Leslie model is 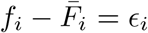 and moreover, 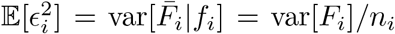.Such demographic deviations mean that the proportions of mothers in each age-class will fluctuate around some stationary distribution. That is, denoting 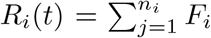 as the total reproduction from age-class *i* and *R*(*t*) as the total reproduction over all age-classes, the observed proportion of mothers of age *i* in the simulation, *R*_*i*_(*t*)*/R*(*t*), will not exactly correspond to *ρ*_*i*_ in the theoretical model. A comprehensive investigation of the effects of population size in simulated processes will provide a further avenue for further research, but is beyond the scope of the present article.

The present work is restricted by assuming a time-invariant, one-sex, age-structured population. Extending to time-dependent vital rates would reflect more realistic population dynamics. The main theoretical challenge here would be to probabilistically sample mothers from a non-stable population (i.e., defining *ρ*_*x,t*_). Methods in Butterick et al. (2025) pertaining to time-inhomogeneous genealogical Markov chains can readily achieve as much. The remainder of moving to time-varying demographics should be merely a case of careful indexing. Incor-porating two sexes should also be relatively straightforward. Model ingredients requiring change would include projecting (i) survival independently by sex and (ii) the numbers of newborns distributed by sex. Constructing block-structured matrices similar to those used in Caswell (2022), with 𝕌^f,m^, 𝔽^f,m^ *in lieu* of the projection matrices therein, would allow such progress. In this case, projecting older and younger lineages (Eq (13) and Eq (16)) would require little change; except for imposing a female dominance in direct ancestor reproduction, and the output being a block-structured vector of female and male kin-number pmfs. Similarly, Focal’s descendants would simply be represented by a block-structured vector of pmfs. One notable change would be that the probable ages at which ancestors reproduce would be sex-specific (i.e., 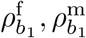 respectively represent the probable ages of Focal’s mother and father at her birth). As such, in order to project direct ancestors, Eq (19) would involve a double summation over both variables (both sexes). We are currently working on extending the present framework to accommodate both sexes, and within a time-variant demography. Good progress has been made here; we hope this development will be the focus of a forthcoming paper.

An interesting but challenging prospect is to extend the present research to a multi-state model. Such progress would require more in-depth consideration. For instance, construction of the matrices in Section 2.4 and Section 2.5 that form the core of this method would become complicated. As well as a function of age, the probability that *j* = 1, 2, … kin die at some given time will also depend on their stages. Elements of the projection matrix 𝕌 (*s*^*′*^, *s, g, f*) under a multi-state model would have to provide the probabilities that so many kin survive from age *s*^*′*^ to *s*, starting from stage *g* and ending in stage *f*. Progress in this direction would allow for a very rich investigation of kinship structures, perhaps paralleling the complexity of simulation. Including stage would allow for example, one to condition fertility profiles (***ψ***) as dependent on elapsed time since last birth (as well as age). Not only would such an innovation provide a more accurate description for human kinship where it is known that time-proximity between births can create a “sibling constellation” (Morosow and Kolk, 2017), but would also be highly relevant in other species which experience postpartum infertility, such as toothed whales (suborder Odontoceti) (Ellis et al., 2024). Implementing stochastic age×stage structured kinship remains an open research problem.

In summary, the methodology proposed in this paper has, for the first time, enabled projecting a probability distribution of the (i) total number of kin and (ii) age-specific distribution of kin, of a typical population member. Having a complete distribution of kin allows for detailed analysis of probabilities of given numbers of relatives present at different parts of the life course, which is important from the point of view of studying the presence of support networks, availability of care, and more.

## 6 Acknowledgements

We would like to thank Hal Caswell for valuable discussions and helpful comments on this project. All authors acknowledge that the research was supported by [Redacted].

## Appendices

### A Supplementary figures

In Figure A-1 we compare the overall kin-number pmfs of total younger and older sisters of Focal predicted by our model (orange) to a stochastic simulation (black), when Focal is aged 40. In Figure A-2 we extend the comparison over all ages of Focal, and compare the probabilities that Focal experiences *j* = 0, 1, 2, 3, 4 sisters.

**Figure A-1:**
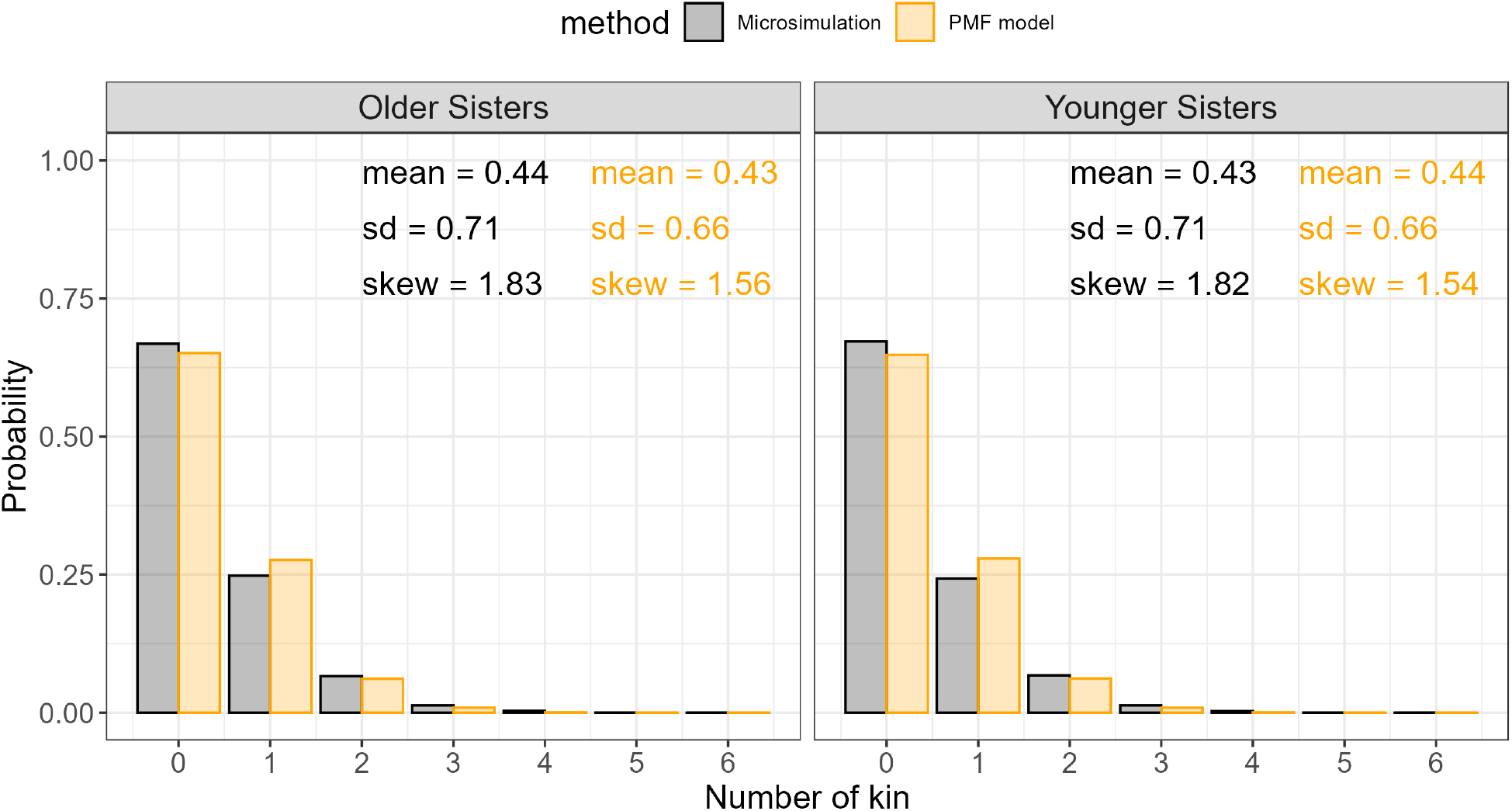
Accumulated-kin pmfs for Focal’s sisters when Focal is aged 40. Legend (and colour) compares the theoretical model to an agent based simulation. Left: older sisters; right: younger sisters. Bars show the probability that Focal has a given number of sisters.

**Figure A-2:**
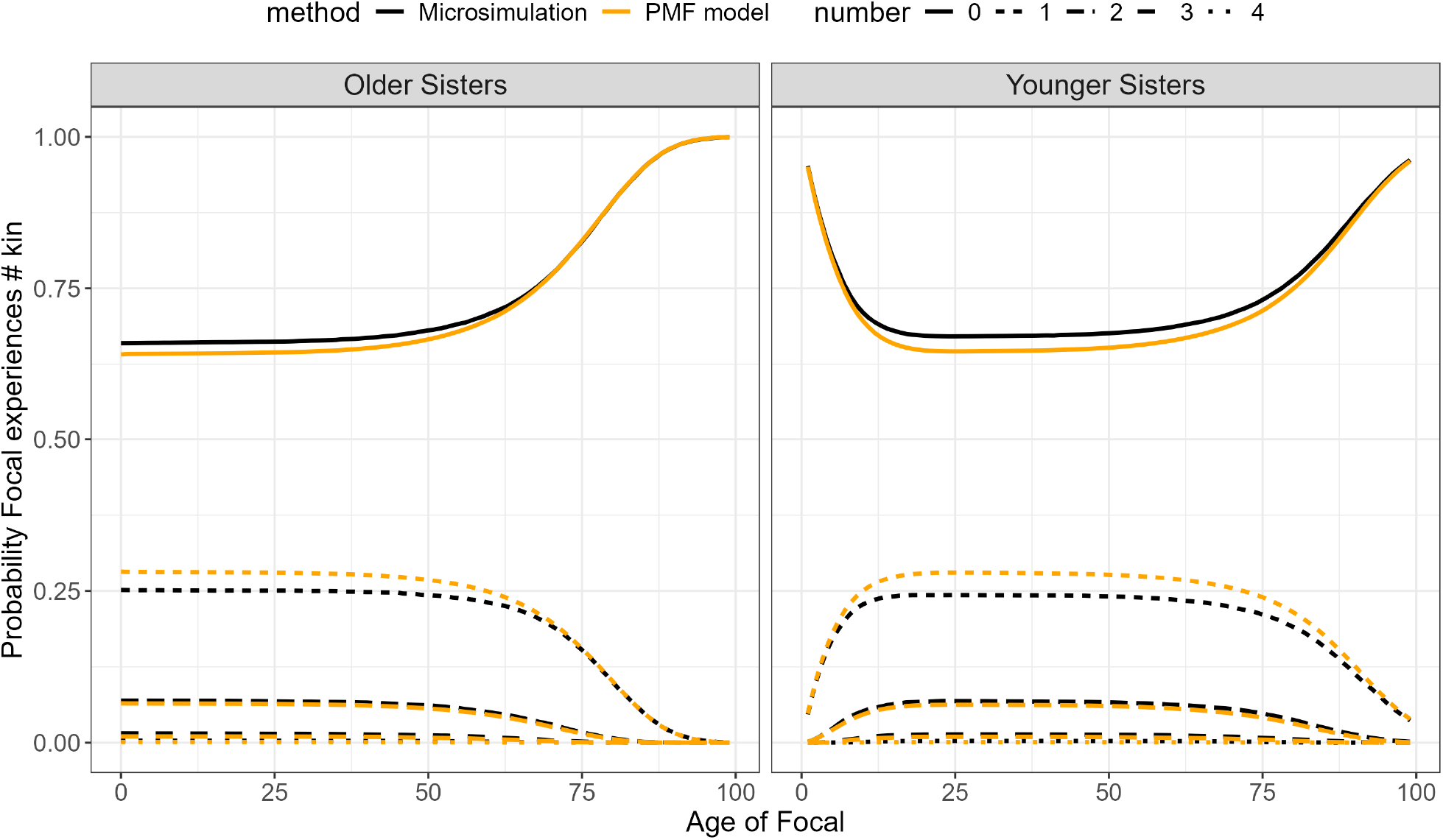
Probable number of sisters over Focal’s lifetime. Legend (and colour) compares the theoretical model to an agent based simulation. Left: older sisters of Focal; right: younger sisters. Line-type shows the probabilities that Focal has a given number of kin.

In Figure A-3 we compare our model’s predicted kin-number pmf for Focal’s mother and Focal’s daughters (orange) to results obtained through a stochastic simulation (black). Shown are pmfs for two ages of Focal. In Figure A-4 we compare, for all ages of Focal, the probability that Focal has *j* = 0, 1, 2, 3, 4 daughters, and the probability that her mother is alive.

**Figure A-3:**
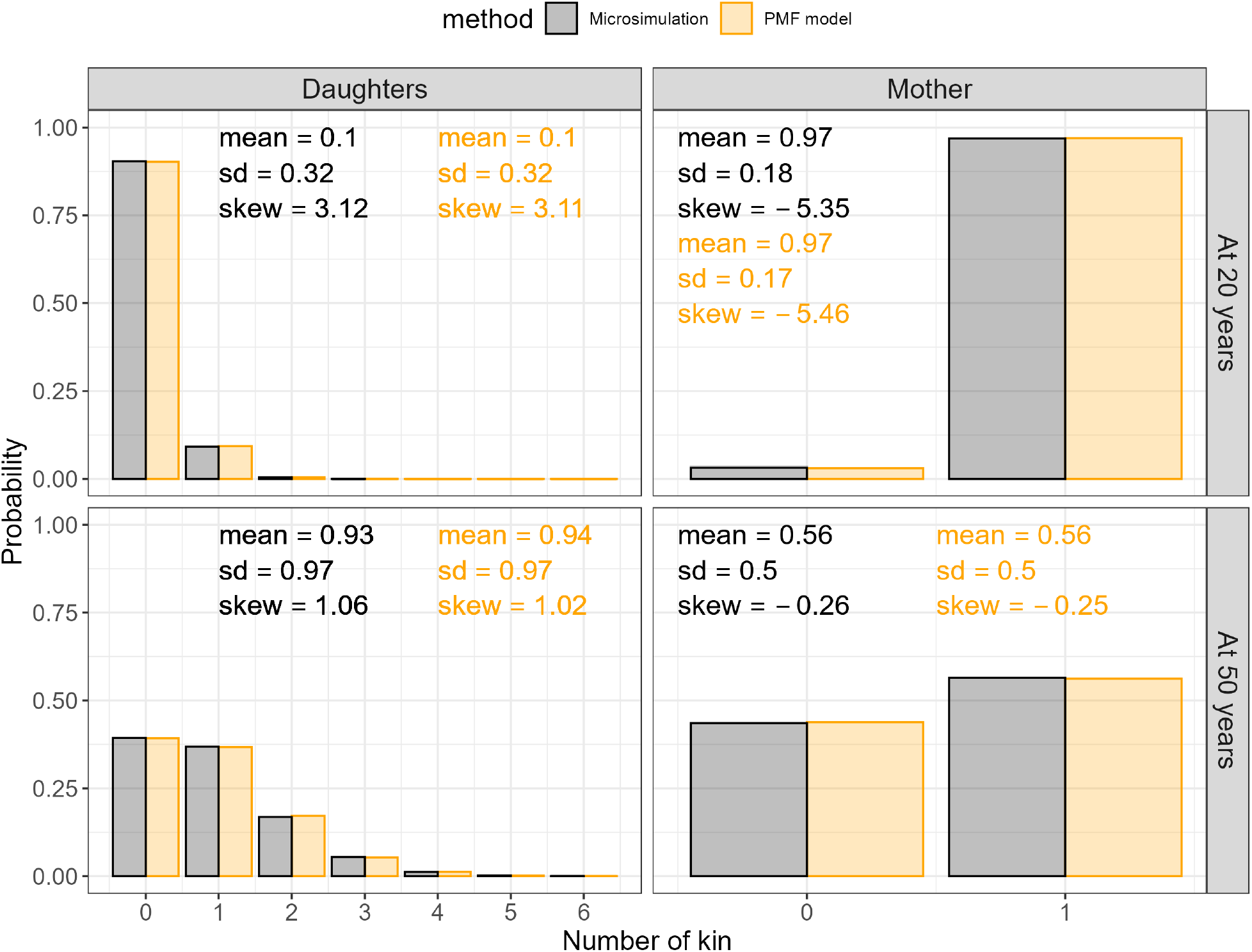
Accumulated-kin pmfs for Focal’s daughters and mother when Focal is aged 20 and 50. Legend (and colour) compares the theoretical model to an agent based simulation. Rows show age of Focal. Left: daughters; right: mother. Bars give the probability that Focal has a given number of kin.

**Figure A-4:**
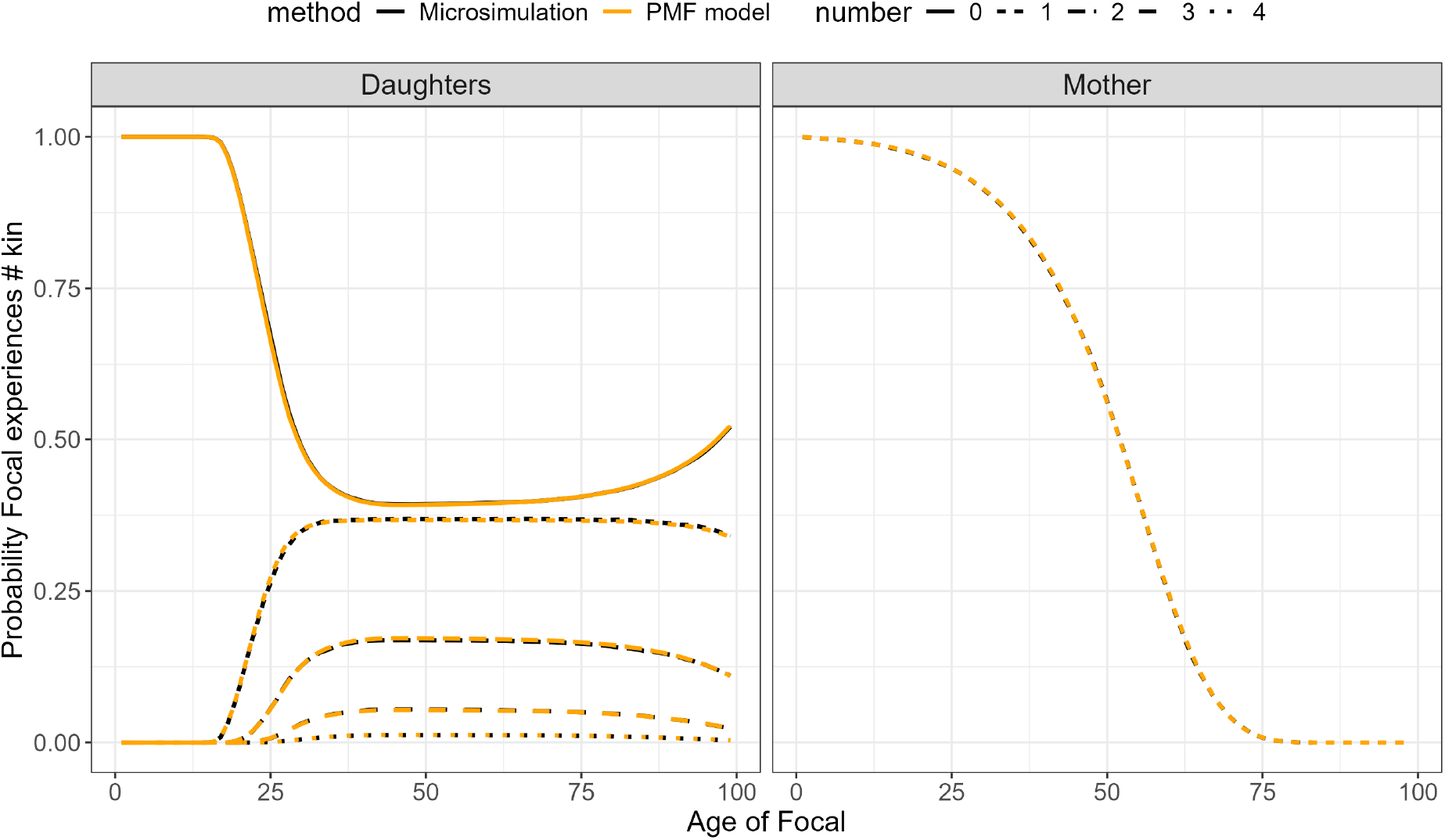
Probable parity of Focal (number of daughters) and the probability her mother is alive, by age of Focal. Legend (and colour) compares the theoretical model to an agent based simulation. Left: Focal’s daughters; right: Focal’s mother. Line-type compares the probabilities that Focal experiences a certain number of kin.

### B The matrix U

The matrix is column-stochastic as can be checked by observing that entries of the *j >* 1 columns obey the binomial expansion:

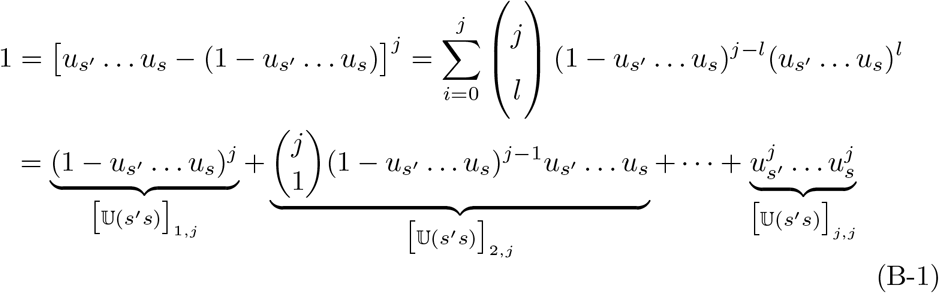

### C Proof of Theorem 1 (and gentle reminder of convolutions)

Before providing the theorem proof we gently remind the reader how a discrete convolution operates on two distributions. The convolution measures how one distribution is augmented by another distribution. As the domain of one distribution is translated over the other, the overlap in the two distributions’ domains is calculated. At each position in the translation, the sum of the element-wise products of the distribution ranges’ is calculated, producing the positional entry in a new distribution.

In the context of kinship, assume that there are two reproductive ages, 15 and 16. Assume that at each age no more than 2 offspring can be born. Let ***ψ***_15_ = (*ψ*_15_(0), *ψ*_15_(1), *ψ*_15_(2))^*†*^ and ***ψ***_16_ = (*ψ*_16_(0), *ψ*_16_(1), *ψ*_16_(2))^*†*^. As ***ψ***_**16**_ is translated over ***ψ***_**15**_, we calculate the convolution schematically:

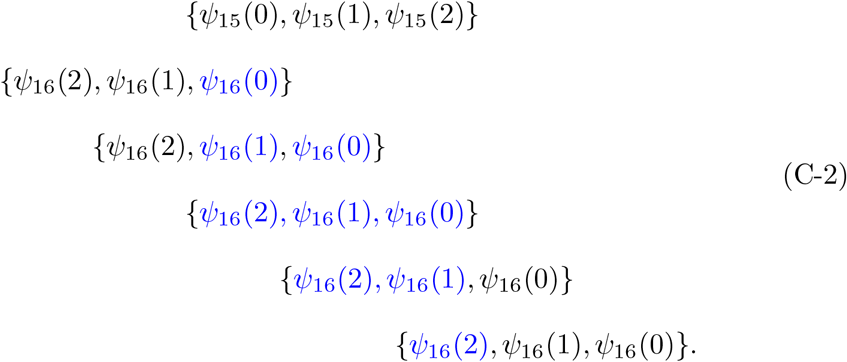

Note that C-2 has ***ψ***_**15**_ as the top row, and each subsequent row consists of ***ψ***_**16**_ shifted rightwards by one element. The first entry of the resulting distribution consists of the blue entry in the second row of C-2 multiplied by the element of ***ψ***_**15**_ that it is vertically aligned with. Similarly, the second entry consists of sum of the product of the two blue elements in the third row of C-2 with the corresponding aligned elements of ***ψ***_**15**_, and so on. The resulting distribution has first entry *ψ*_15_(0)*ψ*_16_(0): the probability that no offspring are born. The second entry is *ψ*_15_(0)*ψ*_16_(1)+*ψ*_15_(1)*ψ*_16_(0): the probability exactly one offspring is born at 15 or 16. The third, fourth and fifth entries respectively yield probabilities that there are exactly 2, 3 and 4 offspring born over the ages 15 and 16. Hence the convolution of the age-specific offspring-number distributions yields the overall offspring-number distribution over “all possible” reproductive ages.

#### C.1 Proof of Theorem 1

*Proof*. Recall that the *j*-th entry of **k**_*s*_(*y*) gives the probability that Focal has *j* − 1 of these kin of age *s* when she is aged *y*, i.e., 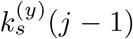.Let 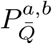 be the probability that Focal has exactly 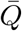 kin at age *y* whereby kin can range between ages *a* and *b*. Suppose that the ages of kin can range only from *a* = *z*_1_ and *b* = *z*_2_ for example. Then

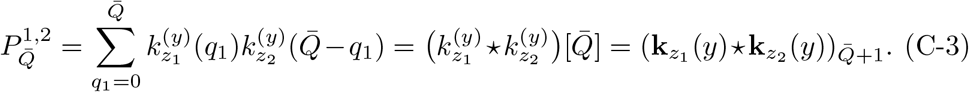

Thus, 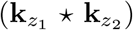 is a discrete probability distribution with *j*-th entry the probability that Focal has exactly *j* − 1 kin when aged *y*. If kin can be of ages *z*_1_, *z*_2_, *z*_3_ then

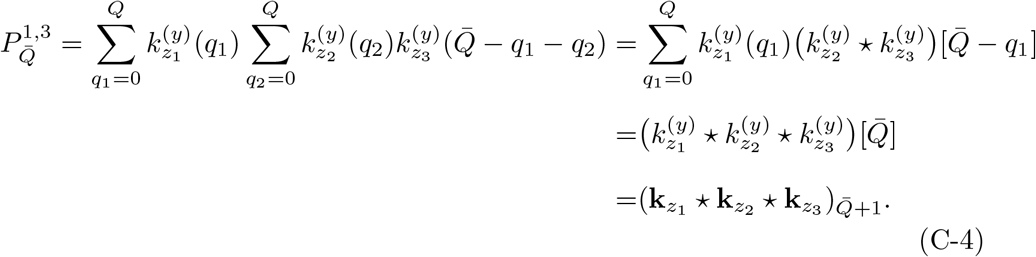

For any age *y* of Focal, the above reasoning leads us to believe that we can extend the convolutions of age-specific pmfs **k**_*z*_(*y*) over all kin potential ages *z*_*i*_, *i* = 1, …, *n* to obtain the number distribution. We use induction to show that if above holds for kin up to age *z*_*m*−1_ then it holds for *z*_*m*_ (for notational ease we omit the superscript *y*). Suppose that the equality holds for *z*_1_ to *z*_*m*−1_, we see that by the induction hypothesis:

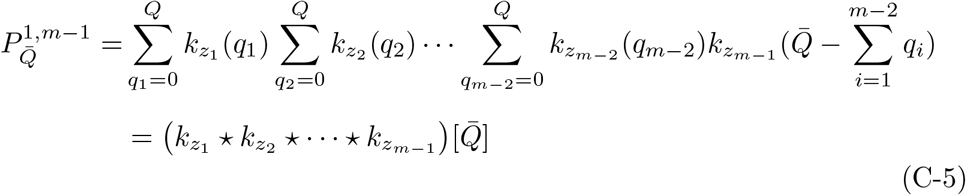

By shifting indices *i* → *i* + 1, by the induction hypothesis

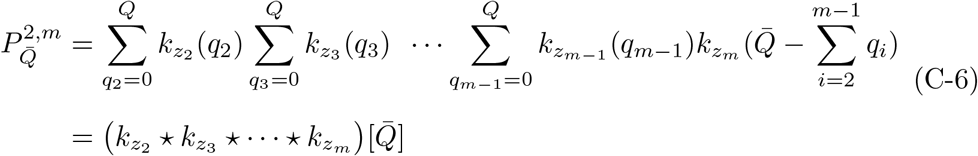

i.e., the 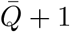-th entry of the distribution 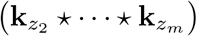 gives the probability that there are 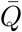 kin over ages *z*_2_ to *z*_*m*_. The probability that there are 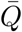 kin over ages *z*_1_ to *z*_*m*_, is therefore given by:

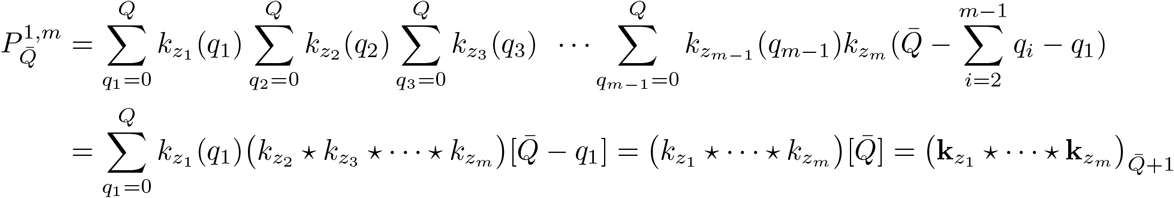

### D Kin which descend through same-age sisters of Focal’s (*q* − 1)-th ancestor

Let the random variable *S* represent the total number of offspring of Focal’s *q*-th ancestor’s, at the age when she had Focal’s (*q* − 1)-th ancestor. Let *S* have distribution ***U***_*s*_. Let the random variable representing the additional “same-age” sisters of Focal’s (*q* − 1)-th ancestor be *A*. Then *A* = *S* − 1. Suppose that *A* has distribution ***ϕ***_*s*_.

By defining

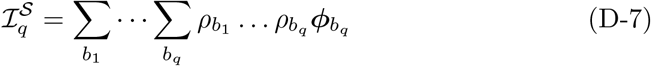

and using the operator

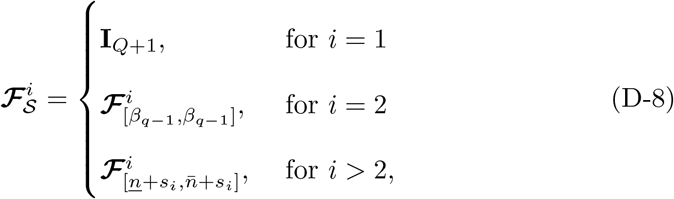

the age-number probability distribution for kin which descend through same-age class sisters of Focal’s (*q* − 1)-th ancestor can be written:

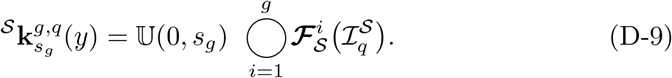

Notice that since *s*_1_ = *β*_*q*−1_, for *i* = 2, the convolution over *s*_1_ reduces to one term:

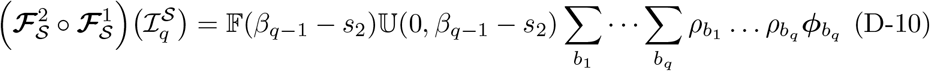

only accounting for reproduction of Focal’s *q*-th ancestor who belong to a group of children born at the same time as Focal’s (*q* − 1)-th ancestor. All subsequent convolutions over reproductive ages extend to the limits *n* and 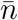.

Within an age-structured demography all offspring are born into the first age-class. Recall that Focal’s *q*-th ancestor produces Focal’s (*q* − 1)-th ancestor at age *b*_*q*_. This observed event means that the reproduction of Focal’s *q*-th ancestor at this age, producing the additional same-age sisters of Focal’s (*q* − 1)-th ancestor, is not independent. We must condition on the given knowledge that one offspring is born: 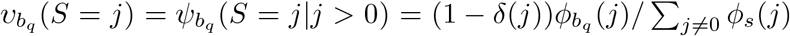.Then using the fact that *S* = *A* + 1, we have 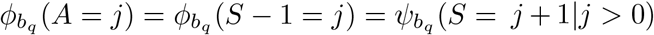.Under the Poisson assumption in Section 4 we recover the so-called zero-truncated Poisson: 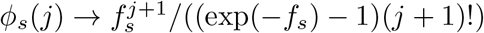

### E Illustration of the formulae in relation to Caswell’s approach

In this section we illustrate our theoretical approach with annotation for ease of interpretation. We also relate our model to the established one of Caswell (2024) by referring to the notation used therein.

#### E.1 Older sisters (m) or (*g* = 1, *q* = 1)-kin

Using notation of the established kinship model (see Figure 1), through Eq (13) our model produces the following pmf for Focal’s older sisters of age *s*_1_, when Focal is aged *y*:

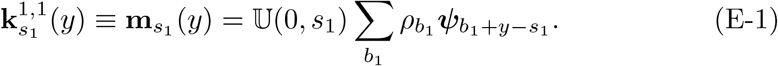

Eq (E-1) is a function of Focal’s age *y* and Focal’s older sister’s age *s*_1_. We interpret the above by conditioning on the age of Focal’s mother at each possible age at which she could have had Focal, *b*_1_. For Focal to experience an older sister of age *s*_1_ when she is age *y*, mother must produce this kin at age *b*_1_ + *y* − *s*_1_. Given that mother has Focal at age *b*_1_ with probability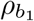, the summation calculates the weighted average reproduction pmf of Focal’s mother “before” Focal is born, as a function of *s*_1_. Pre-multiplication through 𝕌 (0, *s*_1_) calculates the pmf of the offspring who survive to be Focal’s older sisters.

#### E.2 Nieces from older sisters (p) or (*g* = 2, *q* = 1)-kin

Appealing to Eq (13), when Focal is of age *y*, the age-specific pmfs of her nieces aged *s*_2_, born from her older sisters are given through

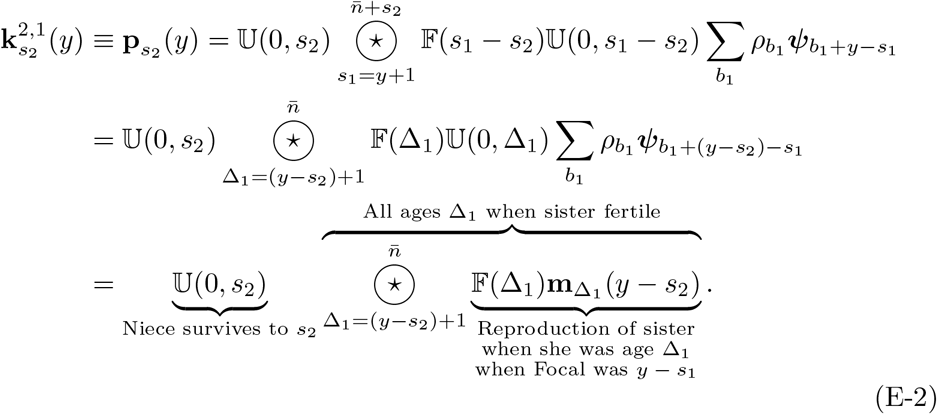

From right to left, Eq (E-2) effectively calculates the complete age-specific pmfs for older sisters of Focal for ages whereby they could potentially produce a niece of Focal, i.e., when Focal was aged *y* − *s*_2_. The convolution over *s*_1_ calculates the total reproduction of Focal’s older sister – yielding a pmf of offspring produced by sister of age *s*_1_ − *s*_2_. The matrix multiplication through 𝕌 (0, *s*_2_) projects the offspring (i.e., newborns nieces of Focal) up to age *s*_2_ when Focal is *y*. The change of variables to Δ_1_ in the last line demonstrates how, as should be expected, nieces of age *s*_2_ of Focal age *y* are produced by Focal’s older sisters at ages Δ_1_ (at least *y* + 1 − *s*_2_) when Focal was age *y* − *s*_2_. Nieces then survive from age 0 up to age *s*_2_.

#### E.3 Aunts older than mother (r) or (*g* = 1, *q* = 2)-kin

Appealing to Eq (13) we obtain the age-specific pmf for aunts who are older than Focal’s mother:

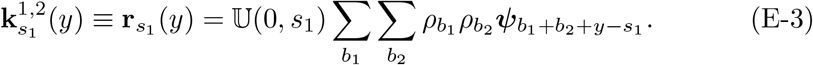

Eq (E-3), through the double summation calculates the weighted average of the pmfs of grandmother’s reproduction at age *b*_1_ + *b*_2_ + *y* − *s*_2_, “before” producing Focal’s mother. The pmf of the newborns, constituting Focal’s aunts older than mother, are then projected through the 𝕌 matrix to survive to age *s*_1_.

#### E.4 Cousins from aunts older than mother (t) or (*g* = 2, *q* = 2)-kin

Appealing to Eq (13) we obtain the age-specific pmf for cousins who descend through aunts who are older than Focal’s mother:

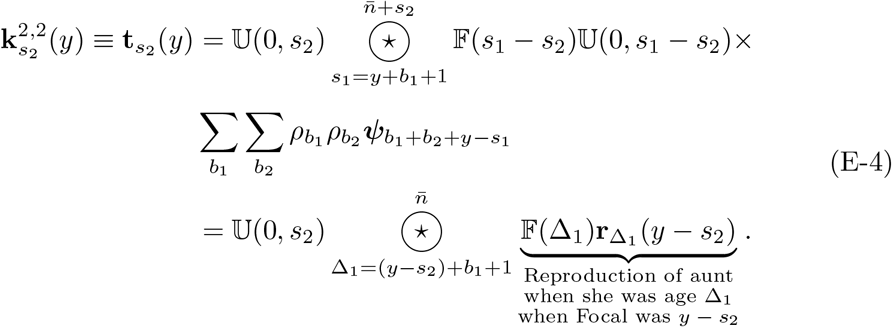

Eq (E-4) reads that cousins of Focal at age *s*_2_ when Focal is *y* were produced by Focal’s older aunts when Focal was *y* − *s*_2_ and when aunt was age Δ_1_ (at least *y* + *b*_1_ + 1 − *s*_2_ or older). In more detail, the double summation calculates the the pmf representing Focal’s grandmother’s reproduction “before” producing Focal’s mother. The convolution over *s*_1_ of the pmf pre-multiplied through 𝔽 (*s*_1_ − *s*_2_) 𝕌 (0, *s*_1_ − *s*_2_) yields the total reproduction of aunts who survive up to and reproduce at age Δ_1_ = *s*_1_ − *s*_2_. Focal’s aunt’s possible age is constrained such she is older than Focal’s mother (i.e., *s*_1_ − *s*_2_ *> y* − *s*_2_ + *b*_1_), explaining the limits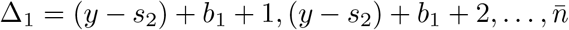. Reproduction of aunt results in the newborn cousins of Focal. Through 𝕌 (0, *s*_2_) the pmf of newborn cousins is projected to age *s*_2_.

#### E.5 Younger sisters (n) or (*q* = 1, *g* = 1)-kin

In relation to the established model Caswell (2019), we have from appealing to Eq (16), the pmf of younger sisters of age *s*_1_ of Focal age *y*:

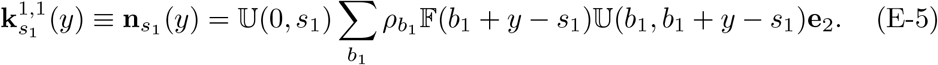

Eq (E-5) reads: at age *b*_1_ Focal’s mother has Focal and at that age her pmf is a unit vector with mass in the 1-kin-number-class (mother is certainly alive when having Focal). Mother survives *y* − *s*_1_ years (through the 𝕌-matrix), up to age *b*_1_ + *y* − *s*_1_, when she produces a younger sister of Focal (through the 𝔽-matrix). The younger sister survives from birth up to age *s*_1_.

#### E.6 Nieces from younger sisters (q) or (*q* = 1, *g* = 2)-kin

We have from Eq (16):

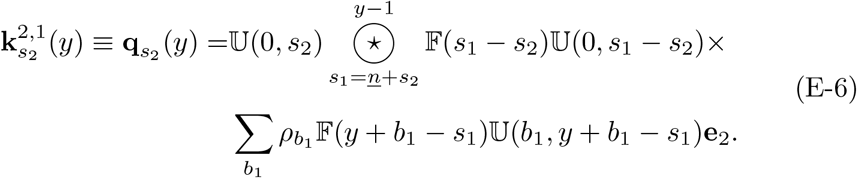

Eq (E-6) reads: at age *b*_1_ Focal’s mother had Focal, and is a unit vector with mass in the 1-kin-number-class (mother is certainly alive when having Focal). Mother then survives up to age *b*_1_ + *y* − *s*_1_ (through the 𝕌-matrix in the summand) at which time she produces a younger sister of Focal (through the 𝔽-matrix). The younger sister survives to age *s*_1_ − *s*_2_, at which point she produces offspring –nieces of Focal, who survive to be age *s*_2_ when Focal is age *y*.

#### E.7 Aunts younger than Focal’s mother (s) or (*q* = 2, *g* = 1)- kin

Appealing to Eq (16) we see that the age-specific pmf for an aunt younger than Focal’s mother who is of age *s*_1_ when Focal is age *y*, is given through

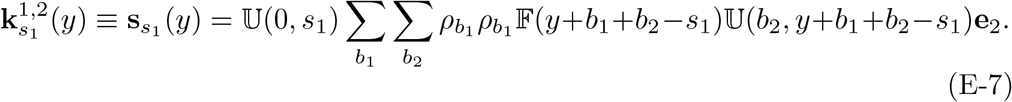

#### E.8 Cousins from aunts younger than Focal’s mother (v) or (*q* = 2, *g* = 1)-kin

We have from Eq (16):

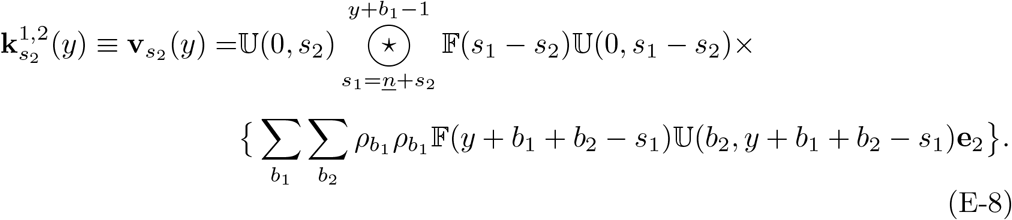

Eq (E-8) reads that cousins of Focal at age *s*_2_ when Focal is *y* were produced by Focal’s younger aunts when Focal was *y* − *s*_2_ and when aunt was age Δ_1_ (at least *y* + *b*_1_ + 1 − *s*_2_ or older).

#### E.9 Daughters (a) or (*g* = 1, *q* = 0)-kin

At each age *i* of Focal’s reproductive life she gives birth to a non-negative integer *j* = 0, 1, …, *Q* of offspring with respective probabilities *ψ*_*i*_(*j*). The pmfs for daughters aged *s*_1_ when Focal is aged *y* are calculated using Eq (18):

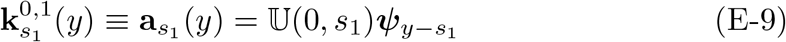

Notice that for Focal’s daughter to be aged *s*_1_ when Focal is aged *y*, Focal’s reproductive probability mass function must have been defined when Focal was at age *y* − *s*_1_, hence 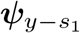

#### E.10 Granddaughters (b) or (*g* = 2, *q* = 0)-kin

We apply Eq (18) with *g* = 2 to obtain the age-specific pmf for granddaughters of age *s*_2_ when Focal is *y*. The equation below demonstrates that the granddaughters of Focal currently aged *s*_2_ are produced through Focal’s daughter over all possible ages of *s*^*′*^ when Focal was age *y* − *s*_2_ (in the past):

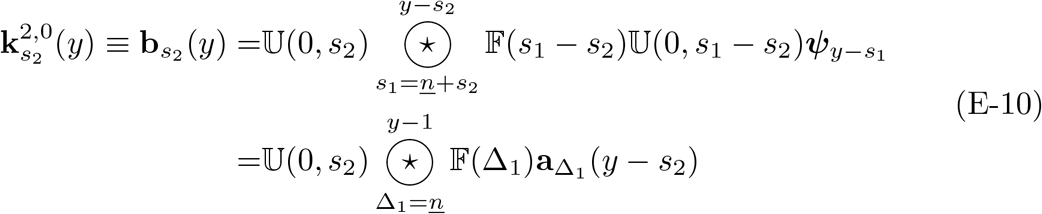

with the change of variables Δ_1_ = *s*_1_ − *s*_2_ rendering Focal’s daughter’s age Δ_1_ at reproduction of Focal’s granddaughter (at which time Focal was age *y* − *s*_2_), and described by the pmf 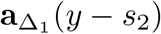.

#### E.11 Mother (d) or (*g* = 0, *q* = 1)-kin

From Eq (19) we see that when Focal is age *y*, the age-specific probability distribution for a mother of age *s*_0_ reduces to

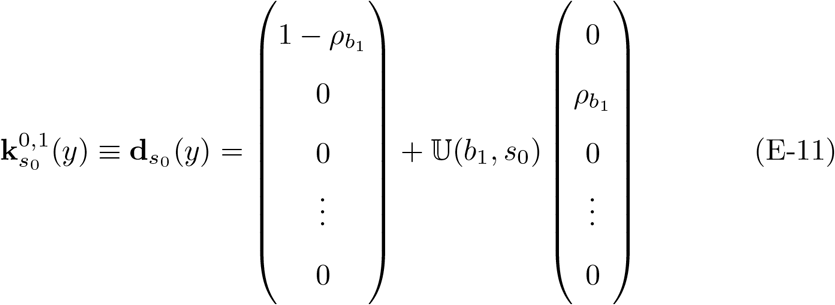

where the summation term in Eq (19) is over one element (i.e., reduced to 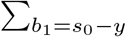 because the age at which mother gives birth to Focal is uniquely defined by mother’s age *s*_0_ when Focal is *y*.

#### E.12 Grandmother (g) or (*g* = 0, *q* = 2)-kin

From Eq (19) we see that when Focal is age *y*, the age-specific probability distribution for a mother of age *s*_0_ is given by:

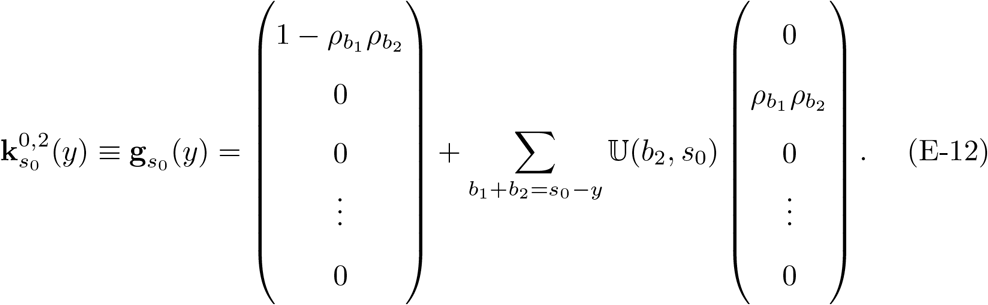

Note that now we are summing over possible ages at which grandmother produced mother (*b*_2_) and mother produced Focal (*b*_1_) which result in grandmother being age *s*_0_ when Focal is *y*.

## Footnotes

1 Based on the data we use here. In other demographies individuals may well experience such numbers of kin.

